# Endothelial cell-derived lactate triggers mesenchymal stem cell histone lactylation to attenuate osteoporosis

**DOI:** 10.1101/2023.03.06.531262

**Authors:** Jinhui Wu, Miao Hu, Heng Jiang, Jun Ma, Chong Xie, Zheng Zhang, Xin Zhou, Jianquan Zhao, Zhengbo Tao, Yichen Meng, Zhuyun Cai, Tengfei Song, Chenglin Zhang, Rui Gao, Hongyuan Song, Yang Gao, Tao Lin, Ce Wang, Xuhui Zhou

## Abstract

Blood vessels play a role in osteogenesis and osteoporosis; however, the role of vascular metabolism is unclear. The present study found that ovariectomized mice exhibit reductions in bone blood vessel density and expression of endothelial glycolytic regulator pyruvate kinase M2 (PKM2). Additional data showed that endothelial cell (EC)-specific deletion of Pkm2 impair osteogenesis and worsen osteoporosis in mice. This was attributed to the impaired differentiation ability toward osteoblast of bone mesenchymal stem cells (BMSCs). Mechanistically, EC-specific deletion of Pkm2 reduce serum lactate levels secreted by ECs, which affect histone lactylation of BMSCs. We identified collagen type I alpha 2 chain, cartilage oligomeric matrix protein, ectonucleotide pyrophosphatase/phosphodiesterase 1, and transcription factor 7 like 2 as histone H3K18 lactylation-regulated osteogenic genes using joint CUT&Tag and RNA-sequencing analyses. The overexpression of PKM2 in ECs, addition of lactate, and exercise were observed to restore the phenotype of endothelial Pkm2-deficient mice. Furthermore, metabolomics of the serum indicated that osteoporosis patients showed a relatively low lactate level. The histone lactylation and related osteogenic genes of BMSCs in osteoporosis patients also decreased. In conclusion, the glycolysis of ECs fuels the differentiation of BMSCs into osteoblasts through histone lactylation, and exercise partially ameliorates osteoporosis through increased serum lactate.

## 1. Introduction

The precise control of osteoblast bone formation and osteoclast bone resorption help to sustain healthy remodeling of the skeleton[1]. Impaired osteoblastic and excessive osteoclastic activity occurs among the elderly and women that have reached menopause, which eventually lead to osteoporosis[2]. Increased bone fragility of osteoporosis easily causes bone fractures, which are associated with increased mortality and healthcare cost[2a, 3]. The differentiation of bone marrow mesenchymal stem/stromal cells (BMSCs) is the major source of osteoblasts, and aging and menopause often disrupt the differentiation of BMSCs into osteoblasts[4]. Blood vessels mediate the transport of oxygen, nutrients, growth factors, and metabolites, which contribute to osteogenesis[5]. Blood vessels, particularly type H vessels affect the differentiation of BMSCs and osteogenesis[6]. However, the role played by vascular endothelial cell metabolism in osteogenesis and osteoporosis is largely unknown.

Vascular endothelial cells (ECs) cover the inner layer of vasculature and regulate the metabolism of other tissues by controlling oxygen and nutrient supply through angiogenesis[7]. In contrast with other cells, ECs produce most of the adenosine triphosphate (ATP) via glycolysis, and angiogenic stimulation further upregulates ECs glycolysis to sustain angiogenesis[8]. ECs prefer aerobic glycolysis regardless of sufficient oxygen in blood and convert most of the glucose to lactate[9]. Large amounts of EC-derived lactate are secreted to the blood, and thus circulated throughout the body. Recent studies indicate that EC-derived lactate regulates the angiogenic phenotype of macrophages/microglia in pathological retinopathy and stimulate the pro-regenerative M2-like phenotype of macrophages to control muscle regeneration[10]. In addition, EC-derived lactate fuels brain pericytes to maintain the blood-brain barrier function[11]. High-intensity exercise at intervals produces large amounts of lactate, which induces cerebral angiogenesis[12]. These studies suggest that EC-derived lactates function as the new signaling molecule in development and pathological conditions.

Lactate-derived lactylation of histone lysine residues serves as epigenetic modification to stimulate gene transcription[13]. Histone lactylation of mouse osteoblast precursor cell line increased during osteoblast differentiation, indicating that lactylation played a role in osteogenesis[14]. Pyruvate kinase catalyzes the final rate-limiting step of glycolysis, thus generating ATP and pyruvate, which is then catalyzed by lactate dehydrogenase A (LDHA) to lactate[15]. Specific deletion of M2 isoform of pyruvate kinase (PKM2) in microglia reduce lactate production and disrupt histone lactylation[16]. ECs express PKM2 exclusively over the M1 isoform of pyruvate kinase (PKM1)[17]. Although the specific deletion of PKM2 in ECs suppresses retinal blood vessel growth, the role played by endothelial PKM2 in skeletal angiogenesis and bone remodeling is largely unknown[17].

The present study investigated the metabolic crosstalk of ECs with BMSCs, and found that ECs use their glycolytic capacity to fuel osteogenesis. In a pathological osteoporosis model (ovariectomy, OVX), both the density of bone blood vessels and expression of PKM2 in ECs decrease significantly. Specific deletion of Pkm2 (Pkm2^ΔEC^) in ECs impairs osteogenesis and worsen the phenotype of osteoporosis in OVX mice. Overexpression of PKM2 in ECs, addition of lactate, and high-intensity interval exercise restored the phenotype of Pkm2-deficient mice. Mechanistically, EC-derived lactate increased the histone lactylation of BMSCs. CUT&Tag and RNA-sequencing analyses reveal that H3K18 lactylation directly stimulates the transcription of osteogenic collagen type I alpha 2 chain (COL1A2), cartilage oligomeric matrix protein (COMP), ectonucleotide pyrophosphatase/phosphodiesterase 1 (ENPP1), and transcription factor 7 like 2 (TCF7L2). Moreover, the metabolomics of the serum indicated that osteoporosis patients have a relatively low lactate level. The histone lactylation and related osteogenic genes of BMSCs in osteoporosis patients also decreased. All the data suggest that EC-derived lactate triggers BMSCs histone lactylation to attenuate osteoporosis.

## 2. Results

### 2.1. Bone blood vessels and endothelial PKM2 decreased in OVX mice

The OVX-induced osteoporosis model is widely used in the study of postmenopausal osteoporosis[18]. The model is constructed as reported in a previous study[19]. Hematoxylin and eosin (H&E) staining analysis showed apparent trabecular bone loss in VOX mice (**Figure S1a**). Microcomputed tomography (micro-CT) further indicated reduced bone mineral density (BMD) and bone volume/total volume (BV/TV) of the distal femur (**Figure S1b-d**). For age-induced osteoporosis, reduced type H vessels (CD 31 and endomucin positive) is associated with osteoporosis; however, total bone blood vessel density is not affected[6a]. Nevertheless, the blood vessel phenotype in OVX mice is elusive. Both total bone blood vessel and type H vessel density decreased in OVX mice (**Figure 1a-c**). ECs are metabolic active cells, and glycolysis is the predominant way of energy production. In this study, we examined the expression of final rate-limiting glycolytic enzyme PKM2 in the bone blood vessel in OVX mice. PKM2 was mainly expressed in the blood vessels of the bone, which might attribute to the highly glycolytic property of vascular endothelial cells (**Figure 1d**). Meanwhile, the results showed significantly decreased expression of PKM2 in bone ECs of OVX mice (**Figure 1d-e**). Western blot analysis of magnet-activated cell sorting (MACS) bone marrow endothelial cells (BMECs) showed apparently attenuated expression of PKM2 in OVX mice (**Figure 1f**). Collectively, these data suggest that endothelial PKM2 may contribute to postmenopausal osteoporosis.

**Figure 1.**
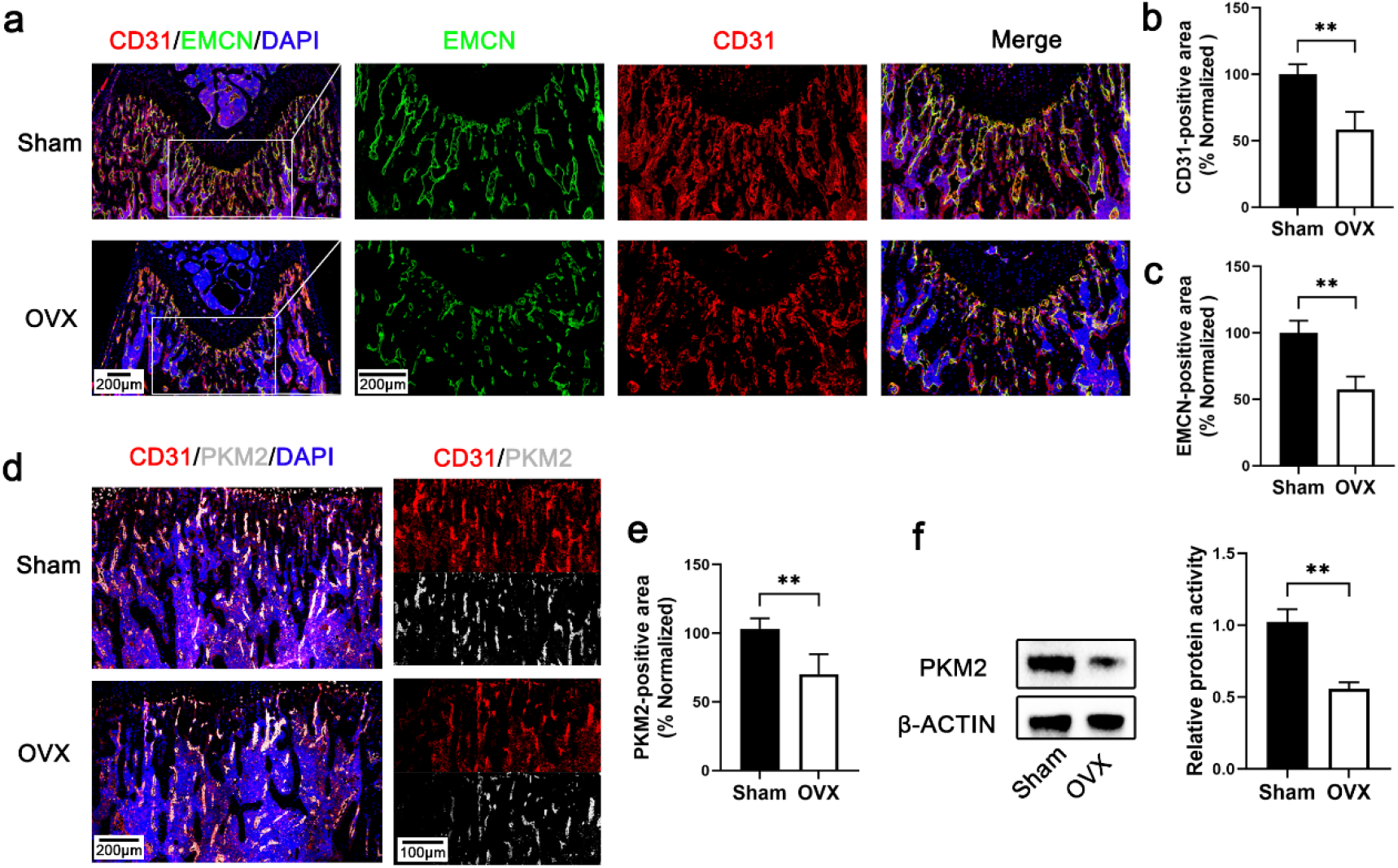
Bone blood vessels and endothelial PKM2 decreased in VOX mice. a) Representative confocal images of sham and VOX mice femurs stained with CD31 (Red) or EMCN (Green) and DAPI (blue) (scale bar 200 μm and 100 μm separately). b) Quantification of CD31 positive vessel area in the BM cavity of the femur sections. n=4, (**p<0.01). c) Quantification of EMCN positive vessel area in the BM cavity of the femur sections. n=4, (**p<0.01). d) Representative confocal images of sham and VOX mice femurs stained with CD31 (Red), PKM2 (white) and DAPI (blue) (scale bar 200 μm and 100 μm separately). e) Quantification of PKM2 positive area in the BM cavity of the femur sections. n=4, (**p<0.01). f) The expression of PKM2 in BMSC sorted from tibiae and femurs of mice. n=3, (**p<0.01).

### 2.2. Endothelial PKM2 controls bone blood vessel density and osteogenesis

To study the role of PKM2 in bone endothelium, Pkm2 ^floxed/floxed^ (Pkm2^fl/fl^) mice were used and intercrossed with Cdh5-Cre^Ert2^ transgenic mice to generate endothelial Pkm2-deleted mice (Pkm2^ΔEC^) (**Figure S2a**). The Pkm2^floxed/floxed^ mice with negative Cdh5-Cre^Ert2^ were used as control (Pkm2^wt^). Western blot analysis of MACS-sorted tibial BMECs showed apparent reduction in the expression of PKM2 in 4-week-old Pkm2^ΔEC^ mice compared to that in control littermates (**Figure S2b-c**). Using Seahorse Flux analysis, we evaluated the glycolytic function of BMECs from Pkm2^ΔEC^ mice (BMECs^Δpkm2^) by measuring the extracellular acidification rate (ECAR). The glycolytic flux in BMECs^Δpkm2^ decreased significantly compared with BMECs from Pkm2^wt^ mice (BMECs^wt^) (**Figure S2d-e**). Immunofluorescence analysis showed that the EC-specific deletion of PKM2 reduced bone blood vessel density significantly in 4-week-old mice (**Figure 2a-b**). Substantially reduced type H vessel density was also observed in Pkm2^ΔEC^ mice (**Figure S2f-g**). Furthermore, the functions of BMECs isolated from the tibiae and femurs of mice were evaluated. The results showed that BMECs^Δpkm2^ exhibited impaired ability of proliferation, migration, and tube formation compared with BMECs^wt^ (**Figure 2c-h**). The data of micro-CT revealed that reduced BMD and BV/TV of the distal femur in 4-week-old Pkm2^ΔEC^ mice compared to those of control littermates (**Figure 2i-k**). These data indicated an important role of endothelial PKM2-mediated glycolysis in osteogenesis.

**Figure 2.**
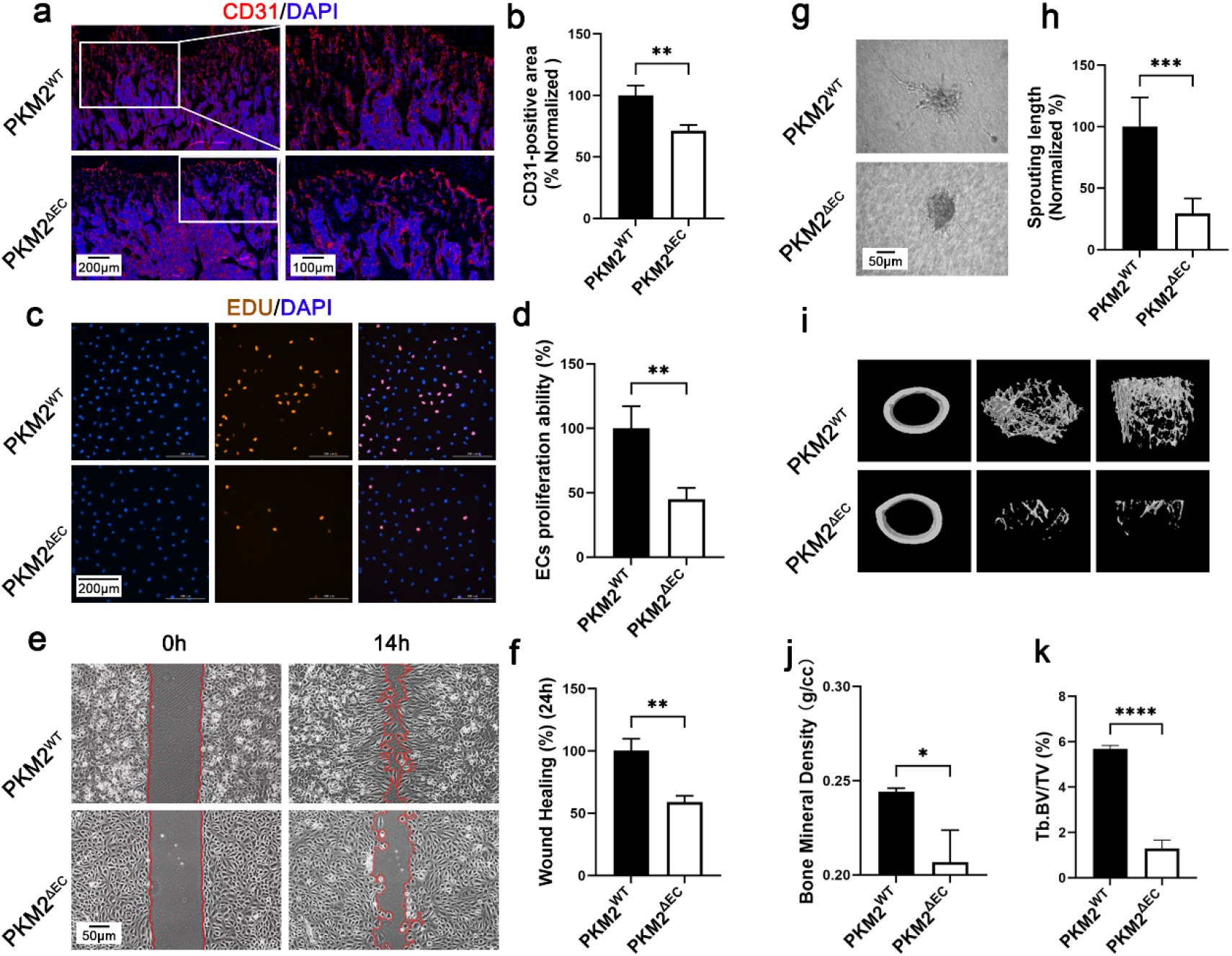
Ablation of endothelial PKM2 decreases bone blood vessels and osteogenesis. a) Representative confocal images of femurs stained with CD31 (Red) and DAPI (blue) (scale bar 200 μm and 100 μm separately). b) Quantification of CD31 positive vessel area in the BM cavity of the femur sections. n=3, (**p<0.01). c) Cell proliferation determined by EdU staining. EdU (Red), DAPI (Blue), (scale bar 200 μm). d) Statistical result of cell proliferation. n=4, (**p<0.01). e) Representative images of migrated cells (scale bar 50 μm). f) Statistical result of cell migration. n=4, (**p<0.01). g) Representative images of cell sprouting (scale bar 50 μm). h) Statistical result of cell sprouting. n=4, (***p<0.001). i) Representative micro-CT images of the distal femur in Pkm2^ΔEC^ mice. j-k) Quantitative analysis of trabecular bone volume per tissue volume (BV/TV) and bone mineral density (BMD) in Pkm2^ΔEC^ mice. n=3, (*p<0.05, ****p < 0.0001).

### 2.3. Deletion of endothelial PKM2 worsens osteoporosis

To study the role of endothelial PKM2 in pathological condition, Pkm2^ΔEC^ mice were used to generate OVX mice. The data indicated that total bone blood vessel and type H vessel density decreased in OVX mice, and EC-specific deletion of PKM2 worsen the phenotype (**Figure 3a-b, Figure S3a-b**). The bone density was evaluated using micro-CT. Compared with ovariectomized Pkm2^wt^ mice, micro-CT analysis showed decreased BMD and BV/TV of the distal femur in ovariectomized Pkm2^ΔEC^ mice (**Figures 3c–e**). Consistent results were acquired using hematoxylin and eosin staining, as the worsened trabecular bone loss was observed in ovariectomized Pkm2^ΔEC^ mice (**Figure S3c**). In addition, the angiogenic properties of BMECs from Pkm2^wt^ mice, ovariectomized Pkm2^wt^ mice, and ovariectomized Pkm2^ΔEC^ mice were evaluated. The data suggested that these BMECs from ovariectomized Pkm2^wt^ mice exhibited impaired ability of proliferation, migration, and tube formation compared to those from Pkm2^wt^ mice (**Figures 3f-k**). BMECs from ovariectomized Pkm2^ΔEC^ mice showed further reduced proliferation, migration, and tube formation ability compared to BMECs sorting from ovariectomized Pkm2^wt^ mice (**Figures 3f-k**). Along with the aforementioned data, the important role of endothelial PKM2 in osteogenesis and osteoporosis was confirmed.

**Figure 3.**
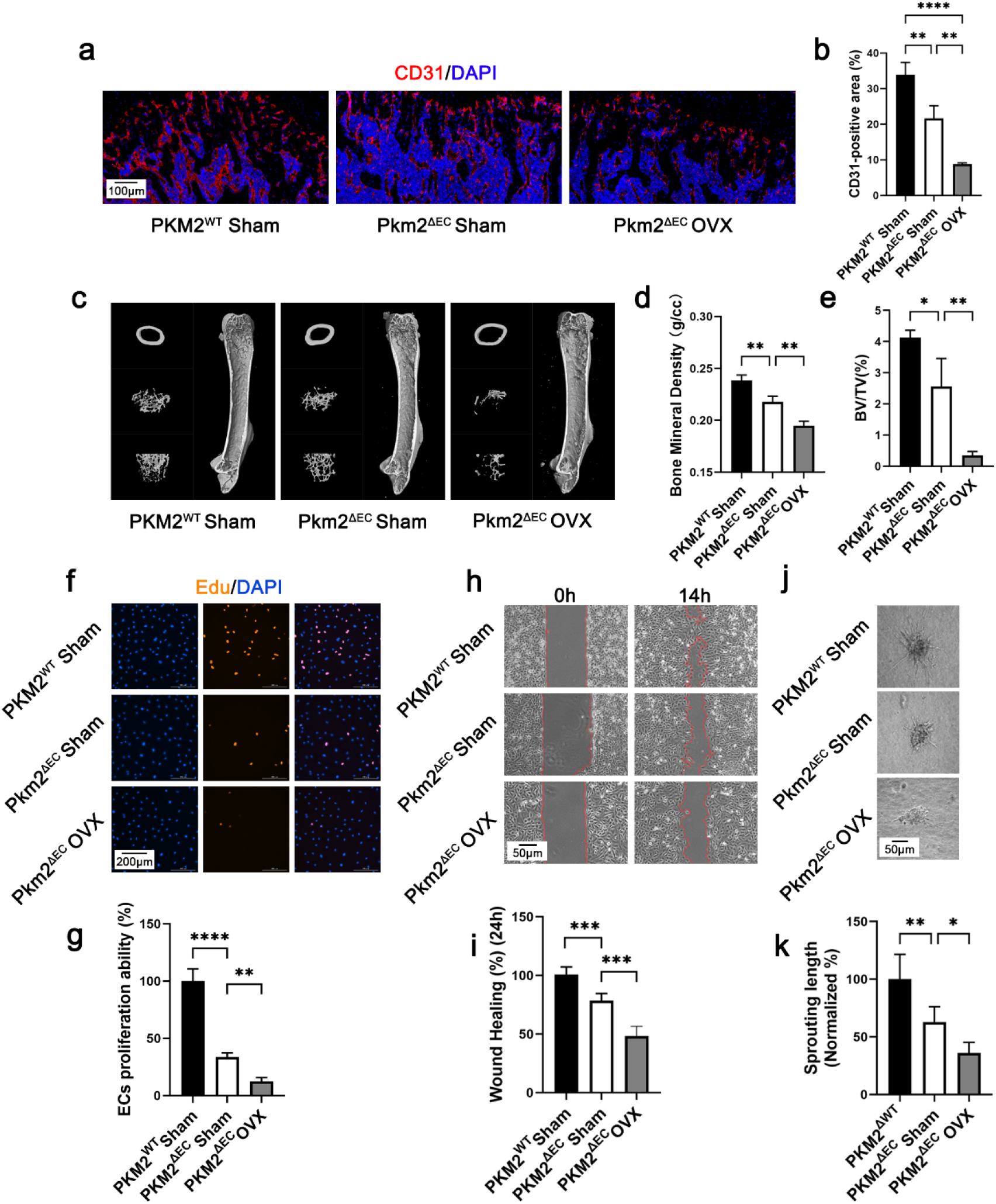
Ablation of endothelial PKM2 worsens osteoporosis. a) Representative confocal images of femurs stained with CD31 (Red) and DAPI (blue) (scale bar 100 μm). b) Quantification of CD31 positive vessel area in the BM cavity of the femur sections. n=5, (**p<0.01). c) Representative micro-CT images of the distal femur in Pkm2^wt^ mice, Pkm2^ΔEC^ mice and OVX Pkm2^ΔEC^ mice. d-e) Quantitative analysis of trabecular bone volume per tissue volume (BV/TV) and bone mineral density (BMD) in Pkm2^wt^ mice, Pkm2^ΔEC^ mice and OVX Pkm2^ΔEC^ mice. n=5, (**p < 0.05, **p < 0.01). f) Cell proliferation determined by EdU staining. EdU (Red), DAPI (Blue), (scale bar 200 μm). g) Statistical result of cell proliferation. n=4, (**p<0.01, ***p<0.001). h) Representative images of migrated cells (scale bar 50 μm). i) Statistical result of cell migration. n=4, (***p<0.001). j) Representative images of cell sprouting (scale bar 50 μm). k) Statistical result of cell sprouting. n=4, (*p<0.05, **p<0.01).

### 2.4. Endothelial lactate affects the differentiation of BMSC to osteoblasts

Recent studies have indicated that bone blood vessels affect osteogenesis through crosstalk with BMSCs[4a, 6a]. To clarify whether the EC-specific deletion of Pkm2 in mice impairs the differentiation of BMSCs, we isolated BMSCs from Pkm2^ΔEC^ mice and Pkm2^wt^ mice. The data showed that BMSCs from Pkm2^ΔEC^ mice exhibited reduced ability to differentiate towards osteoblast using Alizarin Red S (ARS) and Alkaline phosphatase (ALP) staining (**Figures 4a-c**). To confirm the crosstalk between ECs and BMSCs, we sorted BMECs from Pkm2^ΔEC^ mice and Pkm2^wt^ mice and co-cultured them with BMSCs. BMSCs co-cultured with BMECs^Δpkm2^ showed decreased ARS and ALP staining (**Figures 4d-f**). Furthermore, we used PKM2 siRNAs for the knockdown of PKM2 expression in BMECs and co-cultured them with BMSCs. The result was consistent with that of BMECs^Δpkm2^(**Figures S4a-c**). Similar results were acquired using conditional cell culture medium from BMECs^Δpkm2^ and BMECs treated with PKM2 siRNAs (**Figures S4d-i**). PKM2 is a key glycolytic regulator in ECs, and BMECs^Δpkm2^ exhibited suppressed glycolysis flux compared with BMECs^wt^ (**Figure S2b**). The results suggested that EC glycolysis disruption impairs the function of BMSCs; however, the specific factor needs to be clarified.

**Figure 4.**
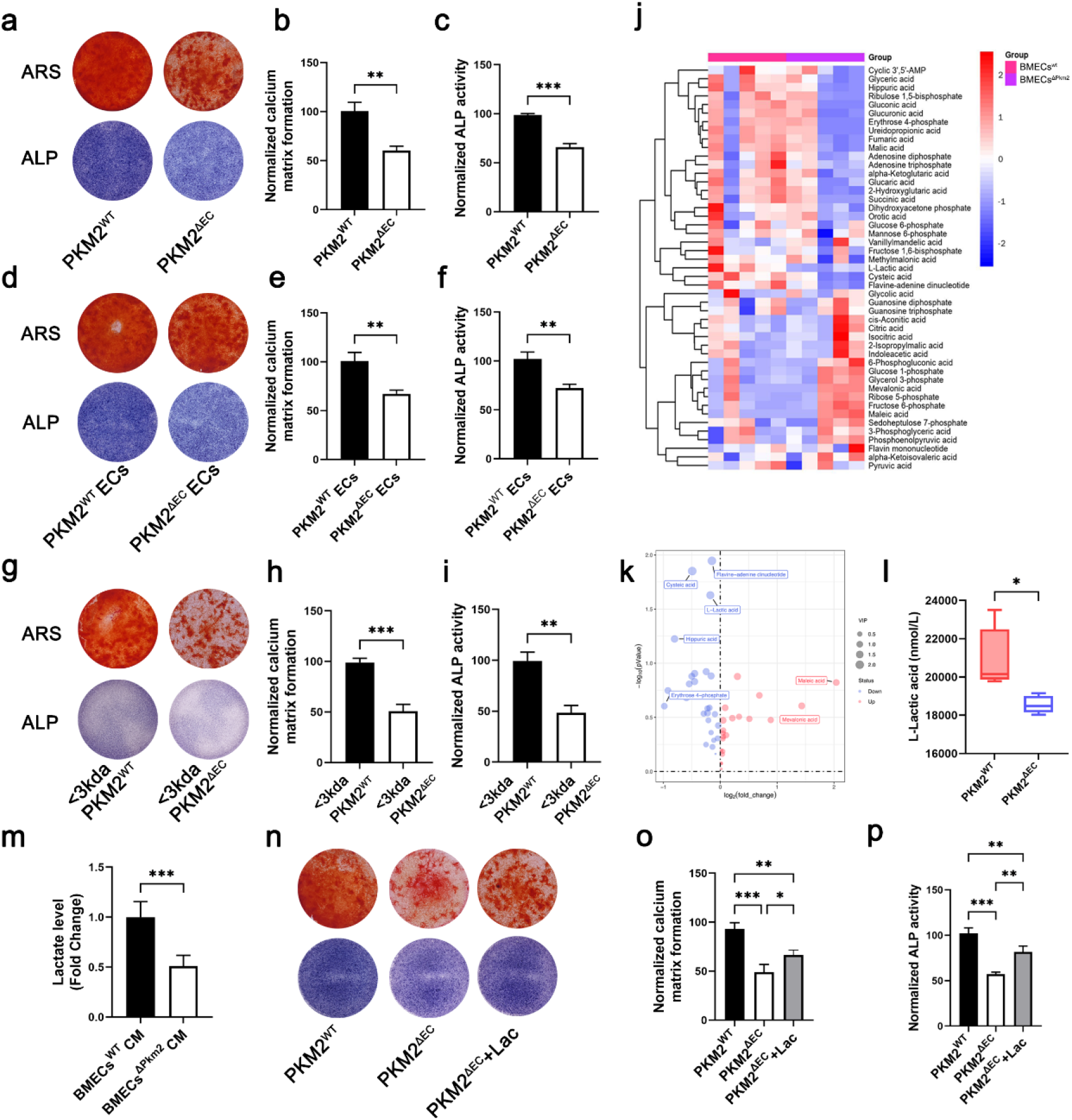
Lactate mediates the crosstalk between BMECs and BMSCs. a) Representative images of ARS and ALP staining of isolated BMSCs from Pkm2^ΔEC^ mice and Pkm2^wt^ mice. b, c) Quantitative analysis of ARS and ALP staining of isolated BMSCs from Pkm2^ΔEC^ mice and Pkm2^wt^ mice. n=4, (**p<0.01, ***p<0.001). d) Representative images of ARS and ALP staining of BMSCs co-cultured with BMECs^Δpkm2^ and BMECs^wt^. e, f) Quantitative analysis of ARS and ALP staining of BMSCs co-cultured with BMECs^Δpkm2^ and BMECs^wt^. n=4, (**p<0.01). g) Representative images of ARS and ALP staining of BMSCs treated with conditional cell culture medium fractions that <3kDa. h, i) Quantitative analysis of ARS and ALP staining of BMSCs treated with conditional cell culture medium fractions that <3kDa. n=4, (**p<0.01, ***p<0.001). j) Heatmap of the metabolites from BMECs^Δpkm2^ and BMECs^wt^ using targeted metabolomics. k) Volcano images of metabolites from BMECs^Δpkm2^ and BMECs^wt^. l) Decreased levels of lactate in BMECs^Δpkm2^ (*p<0.05). m) Decreased levels of lactate in the conditional medium of BMECs^Δpkm2^ n=4, (***p<0.001). n) Representative images of ARS and ALP staining of BMSCs treated with lactate. o, p) Quantitative analysis of ARS and ALP staining of isolated BMSCs treated with lactate. n=4, (*p<0.05, **p<0.01, ***p<0.001).

To better identify the soluble factors in conditional medium, 3K centrifugal filters were used to fractionate conditional cell culture medium according to size (<3 and >3kDa), and both fractions were tested for their activity. The result showed that BMSCs treated with <3kDa fraction from BMECs^Δpkm2^ medium exhibited reduced ARS and ALP staining (**Figures 4g-i**). However, there was no differences between BMSCs treated with >3kDa fraction from BMECs^Δpkm2^ medium and from BMECs^wt^ medium (**Figures S5a-c**). Meanwhile, the boiling of conditional medium did not change its effect on the differentiation of BMSCs (**Figures S5d-f**). These results suggest that the possible factor mediate BMECs crosstalk with BMSCs may be low molecular weight metabolite. ECs exploit glycolysis to generate most of their energy, and lactate is the major metabolite of glycolysis[9]. Targeted metabolomics of BMECs^Δpkm2^ and BMECs^wt^ revealed that intracellular levels of lactate decreased significantly in BMECs^Δpkm2^ (**Figures 4j-l**). Therefore, we tested whether lactate affects BMSCs differentiation. We found that lactate levels were lower in conditional medium derived from BMECs^Δpkm2^(**Figure 4m**). Furthermore, the addition of 5 mM lactate partially rescues the reduced ARS and ALP staining of BMSCs co-cultured with BMECs^Δpkm2^ (**Figures 4n-p**). Collectively, these data indicate that lactate mediate the crosstalk between BMECs and BMSCs.

### 2.5. BMSCs histone H3K18la lactylation boosts osteogenic genes

Lactate-derived lactylation of histone lysine residues directly regulates gene expression[13a]. Therefore, the lactylation of BMSCs from Pkm2^ΔEC^ mice and Pkm2^wt^ mice were evaluated. The results showed that suppressed histone lactylation was detected in BMSCs from Pkm2^ΔEC^ mice (**Figure 5a**). Additional data indicated that H3K18la histone lactylation was affected (**Figure 5a**). To identify candidate target genes regulated by H3K18la histone lactylation, we performed CUT&Tag assay using H3K18la antibody. The result showed that H3K18la was enriched in the promoter and upstream gene regions (**Figure 5b, Figure S6a**). There were 308 upregulated genes and 35 downregulated genes with differential H3K18la modification in BMSCs from Pkm2^wt^ mice (**Figure 5c**). KEGG analysis showed that H3K18la-modified genes were involved in development, energy metabolism, endocrine system and lipid metabolism (**Figure S6b**). Furthermore, Gene Set Enrichment Analysis (GSEA) of H3K18la bound genes showed significant enrichment in “osteoblast differentiation” and “bone development” (**Figure 5d**). To clarify the target genes of H3K18la, we further performed RNA sequencing of BMSCs isolated from Pkm2^ΔEC^ mice and Pkm2^wt^ mice. There were 3949 upregulated genes and 2114 downregulated genes in BMSCs from Pkm2^wt^ mice (**Figure S6c**). Gene ontology (GO) analysis showed that ossification belonged to the top 20 affected categories (**Figure 5e**). Joint CUT&Tag and RNA-seq analyses revealed that there were 58 upregulated genes in BMSCs from Pkm2^wt^ mice (**Figure 5f-g, Figure S6d**). These genes were involved in bone morphogenesis, ossification and bone mineralization (**Figure 5h**). COL1A2, COMP, ENPP1 and TCF7L2 were identified as potential target genes of H3K18la lactylation (**Figure 5g**). A genomic snapshot revealed the H3K18la modification sites in COL1A2, COMP, ENPP1 and TCF7L2 (**Figure S6e**). ChIP-qPCR and RT-qPCR confirm the regulation of H3K18la lactylation on COL1A2, COMP, ENPP1, and TCF7L2 (**Figures 5i, j**). Here, we identified target genes of H3K18la in BMSCs which contributed to osteoblast differentiation.

**Figure 5.**
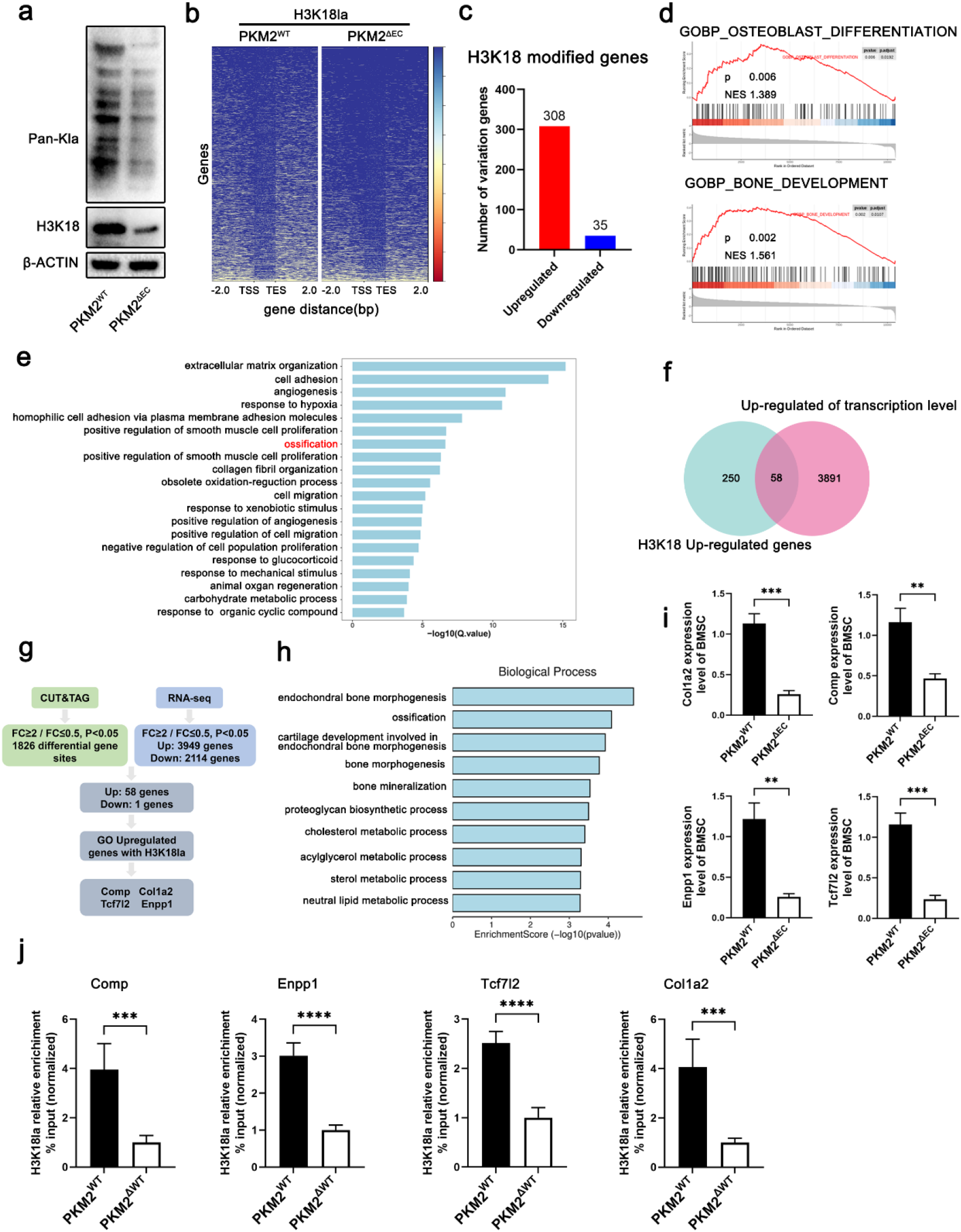
H3K18la lactylation boosts osteogenic genes in BMSCs. a) The expression of pan-Kla and H3K18la decreased in BMSCs isolated from Pkm2^ΔEC^ mice. b) Heatmap for H3K18la binding peaks in BMSCs isolated from Pkm2^ΔEC^ and Pkm2^wt^ mice. c) H3K18la modified genes that differently regulated in BMSCs isolated from Pkm2^ΔEC^ and Pkm2^wt^ mice. d) GSEA of H3K18la bound genes in BMSCs isolated from Pkm2^ΔEC^ and Pkm2^wt^ mice. e) Gene ontology analysis of top 20 ontology terms. f-g) Joint CUT&Tag and RNA-seq analysis filtered COL1A2, COMP, ENPP1 and TCF7L2 as downstream targets of H3K18la. h) Joint CUT&Tag and RNA-seq analysis showed bone morphogenesis, ossification and bone mineralization terms were mainly affected. i) qPCR analysis of COL1A2, COMP, ENPP1 and TCF7L2 in BMSCs isolated from Pkm2^ΔEC^ and Pkm2^wt^ mice. n=4, (***p<0.001, ****p<0.0001). j) ChIP-qPCR analysis of H3K18la binding to the promoter regions of COL1A2, COMP, ENPP1 and TCF7L2 in BMSCs isolated from Pkm2^ΔEC^ and Pkm2^wt^ mice. n=4, (**p<0.01, ***p<0.001).

### 2.6. Endothelial lactate controls differentiation of BMSCs to osteoblasts through H3K18la lactylation

To understand the role of BMSCs H3K18la lactylation in osteogenesis, we generated vascular targeting PKM2 adenovirus to rescue the expression of endothelial PKM2 *in vitro*. The serotype 9 adeno associated virus (AAV9) was used to carry Pkm2 cDNA. The efficiency of PKM2 adenovirus was confirmed using western blot (**Figure S7a**). Overexpression of PKM2 in BMECs^Δpkm2^ rescued the reduced cell proliferation, migration, and tube formation (**Figures S7b-g**). Meanwhile, the decreased glycolytic flux and lactate levels in BMECs^Δpkm2^ were rescued by PKM2 adenovirus (**Figures 6a**,**b**). Then, BMECs^Δpkm2^ infected with PKM2 adenovirus were co-cultured with BMSCs, which improved the decreased ARS and ALP staining in the BMECs^Δpkm2^ co-cultured group (**Figures 6c-e**). H3K18la lactylation and target genes were then evaluated. Overexpression of PKM2 in BMECs^Δpkm2^ increased histone H3K18la lactylation in BMSCs when they were co-cultured (**Figure 6f**). Consistent results were acquired in the expression of target genes (**Figure 6g**). Conditional cell culture medium from BMECs^Δpkm2^ infected with PKM2 adenovirus increased histone H3K18la lactylation and target gene expression compared with medium from BMECs^Δpkm2^ (**Figures 6h, i**).

**Figure 6.**
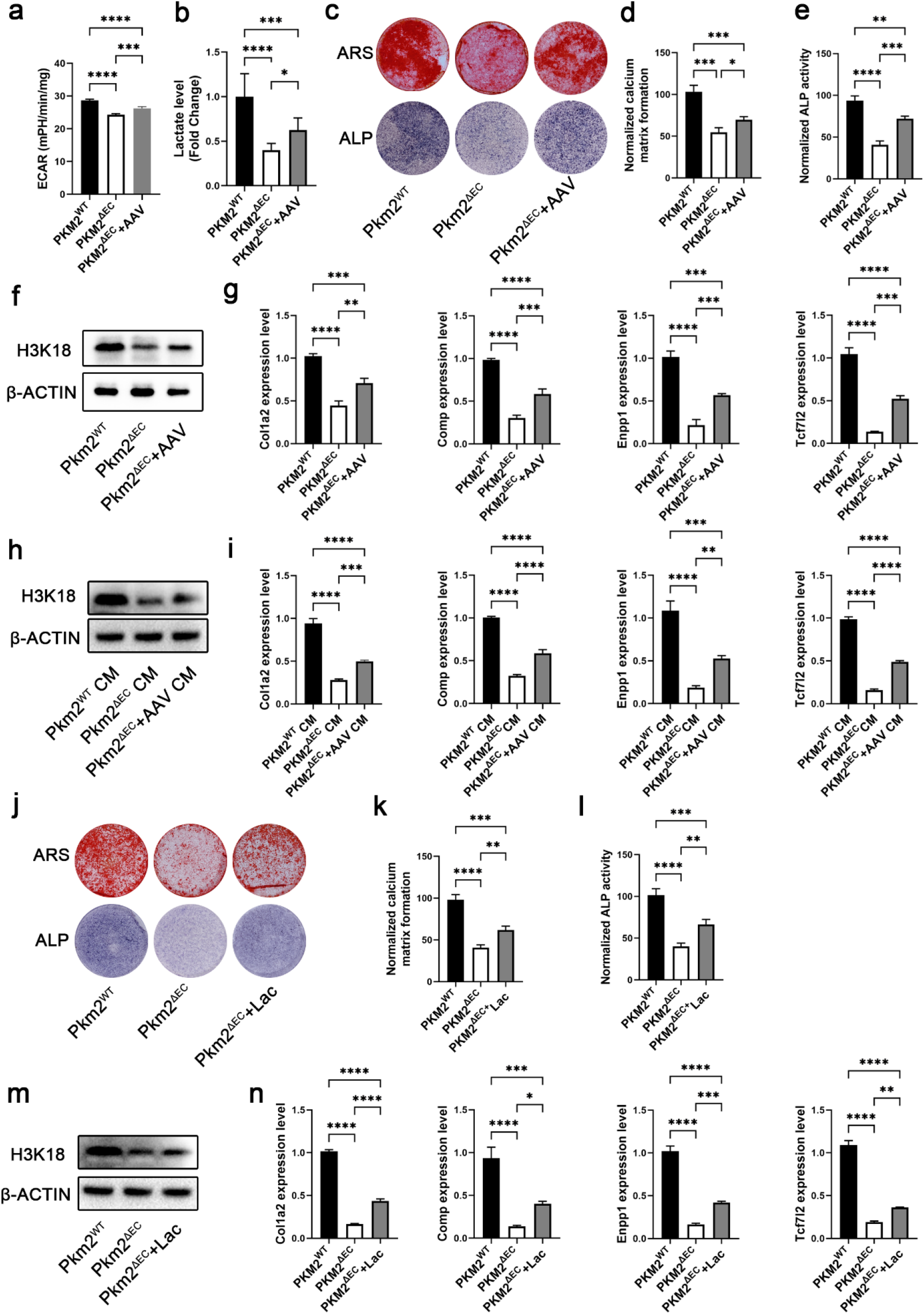
Endothelial lactate contributes to BMSC differentiation towards osteoblasts. a) Overexpression of PKM2 in ECs increases the glycolytic flux. n=3, (***p<0.001, ****p<0.0001). b) Overexpression of PKM2 in ECs increases the production of lactate. n=3, (*p<0.05, ***p<0.001, ****p<0.0001). c) Representative images of ARS and ALP staining of BMSCs co-cultured with ECs that infected with PKM2 adenovirus. d, e) Quantitative analysis of ARS and ALP staining of BMSCs co-cultured with ECs infected with PKM2 adenovirus. n=4, (*p<0.05, **p<0.01, ***p<0.001, ****p<0.0001). f) Histone H3K18la lactylation increased in BMSCs co-cultured with ECs infected with PKM2 adenovirus. g) qPCR analysis of COL1A2, COMP, ENPP1, and TCF7L2 in BMSCs co-cultured with ECs infected with PKM2 adenovirus. n=3, (**p<0.01, ***p<0.001, ****p<0.0001). h) Histone H3K18la lactylation in BMSCs cultured with medium from ECs infected with PKM2 adenovirus. i) qPCR analysis of COL1A2, COMP, ENPP1, and TCF7L2 in BMSCs cultured with medium from ECs infected with PKM2 adenovirus. n=3, (**p<0.01, ***p<0.001, ****p<0.0001). j) Representative images of ARS and ALP staining of BMSCs cultured in conditional medium supplement with lactate. k, l) Quantitative analysis of ARS and ALP staining of BMSCs cultured in conditional medium supplement with lactate. n=4, (**p<0.01, ***p<0.001, ****p<0.0001). m) Histone H3K18la lactylation increased in BMSCs cultured in conditional medium supplement with lactate. n) qPCR analysis of COL1A2, COMP, ENPP1 and TCF7L2 in BMSCs cultured in conditional medium supplement with lactate. n=4, (*p<0.05, **p<0.01, ***p<0.001, ****p<0.0001).

Then, we evaluated the effect of lactate on BMSCs to determine whether it could rescue the phenotype *in vitro*. The addition of 5 mM lactate to the medium of BMSCs from Pkm2^ΔEC^ mice rescued the decreased ARS and ALP staining (**Figures 6j-l**), and upregulated histone H3K18la lactylation (**Figure 6m**). Similar results were acquired in the expression of H3K18la target genes (**Figure 6n**). Furthermore, the addition of lactate to the conditional cell culture medium from BMECs^Δpkm2^ increased the ARS and ALP staining of BMSCs (**Figure S8a-c**).

Meanwhile, histone H3K18la lactylation and its target genes in BMSCs were rescued (**Figure S8d, e**). In summary, restoring EC glycolysis or lactate levels rescue differentiation of BMSCs to osteoblast.

### 2.7. Increasing lactate levels improve bone mineral density in mice

To reveal the metabolic crosstalk between ECs and BMSCs, we explored the effect of histone H3K18la lactylation on osteogenesis *in vivo*. The bone vessel intensity of pkm2^ΔEC^ mice treated with PKM2 adenovirus increased compared with that of pkm2^ΔEC^ mice (**Figure 7a, b**). The lactate levels were evaluated and showed that PKM2 adenovirus treatment increased serum lactate in pkm2^ΔEC^ mice (**Figure 7c**). The micro-CT data revealed that improved BMD and BV/TV of the distal femur in 4-week-old Pkm2^ΔEC^ mice treated with PKM2 adenovirus (**Figures 7d-f**). Similar results were acquired using H&E staining (**Figure S9a**). The osteoporosis was partially rescued by PKM2 adenovirus in the bone of pkm2^ΔEC^ mice as assessed using micro-CT and H&E staining (**Figures 7g-i and S9b**).

**Figure 7.**
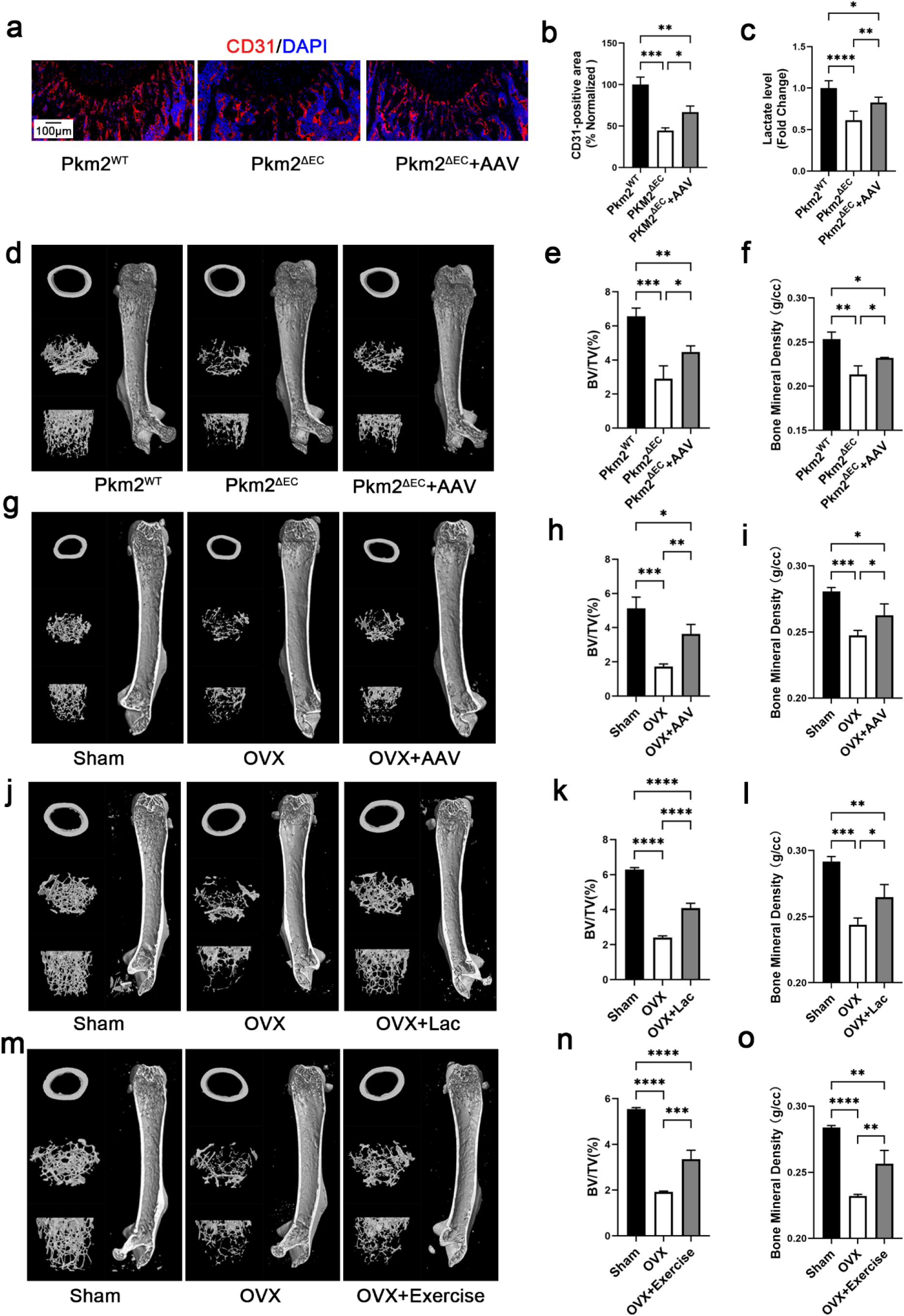
Endothelial lactate restores the phenotype of Pkm2-deficient mice and improves osteoporosis. a) Representative confocal images of femurs stained with CD31 (Red) and DAPI (blue) of Pkm2^wt^ mice, Pkm2^ΔEC^ mice, and Pkm2^ΔEC^ mice treated with PKM2 adenovirus. Scale bars, 100 μ m. b) Quantification of CD31 positive vessel area in the BM cavity of the femur sections. n=4, (*p<0.05, **p<0.01, ***p<0.001). c) PKM2 adenovirus treatment increased serum lactate in pkm2^ΔEC^ mice. n=4, (*p<0.05, **p<0.01, ****p<0.0001). d) Representative micro-CT images of the distal femur in Pkm2^wt^ mice, Pkm2^ΔEC^ mice and Pkm2^ΔEC^ mice treated with PKM2 adenovirus. e, f) Quantitative analysis of trabecular bone volume per tissue volume (BV/TV) and bone mineral density (BMD). n=5, (*p<0.05, **p<0.01, ***p<0.001). g) Representative micro-CT images of the distal femur in Sham, OVX, and OVX mice treated with PKM2 adenovirus. h, i) Quantitative analysis of trabecular bone volume per tissue volume (BV/TV) and bone mineral density (BMD). n=4, (*p<0.05, **p<0.01, ***p<0.001). j) Representative micro-CT images of the distal femur in Sham, OVX and OVX mice treated with lactate. k, l) Quantitative analysis of trabecular bone volume per tissue volume (BV/TV), and bone mineral density (BMD). n=4, (*p<0.05, **p<0.01, ***p<0.001, ****p<0.0001). m) Representative micro-CT images of the distal femur in Sham, OVX, and exercised OVX mice. n, o) Quantitative analysis of trabecular bone volume per tissue volume (BV/TV) and bone mineral density (BMD). n=4, (**p<0.01, ***p<0.001, ****p<0.0001).

Next, we studied whether the addition of lactate could improve osteoporosis in the OVX mice model. The mice were subcutaneously injected with lactate every two days after ovariectomy. Two months later, the bone density was evaluated. The data of micro-CT revealed that BMD and BV/TV of the distal femur were partially rescued after treatment with lactate (**Figure 7j-l**). H&E staining analysis also showed less trabecular bone loss in OVX mice treated with lactate (**Figure S10a**). These data highlight the important role played by lactate in the crosstalk between ECs and BMSCs.

Exercise has been widely recognized as a therapeutic option to prevent osteoporosis; however, the underlying mechanism is still elusive[20]. The present study indicates that exercise-induced lactate may be beneficial. Our data showed that high-intensity interval exercise increased blood lactate levels in OVX mice (**Figure S10b**). The micro-CT data revealed that BMD and BV/TV of the distal femur were partially rescued in OVX mice using exercise (**Figures 7m-o**). Similar results were obtained using H&E staining (**Figure S10c**).

### 2.8. Lactate levels were negatively correlated with osteoporosis through BMSCs histone lactylation

To better reveal the possible role of lactate in osteoporosis, we determined metabolites of glucose metabolism using targeted metabolomics. We collected 86 serum samples from patients with or without osteoporosis according to bone mineral density (BMD) measured using dual-energy X-ray absorptiometry. Age, body mass index, fasting glucose and BMD were summarized in supplementary **Table 1**. The metabolomics analysis showed that 2 metabolites in the osteoporosis group were significantly changed (**Figure 8a-b**). Lactate decreased in the serum of osteoporosis patients (**Figure 8c**). The data suggested that lactate levels may be a negative biomarker for osteoporosis diagnosis and management. To better understand the role of lactate in osteoporosis, we isolated BMSCs from BM aspirates of osteoporosis patients and evaluated the histone lactylation of human BMSCs. The data showed that decreased histone H3K18la lactylation was observed in BMSCs isolated from osteoporosis patients (**Figures 8d-f**). Target genes of H3K18la lactylation were further assessed using qPCR, which indicated that the expression of osteogenic genes, COL1A2, COMP, ENPP1, and TCF7L2, were decreased in BMSCs isolated from osteoporosis patients (**Figures 8g**).

**Figure 8.**
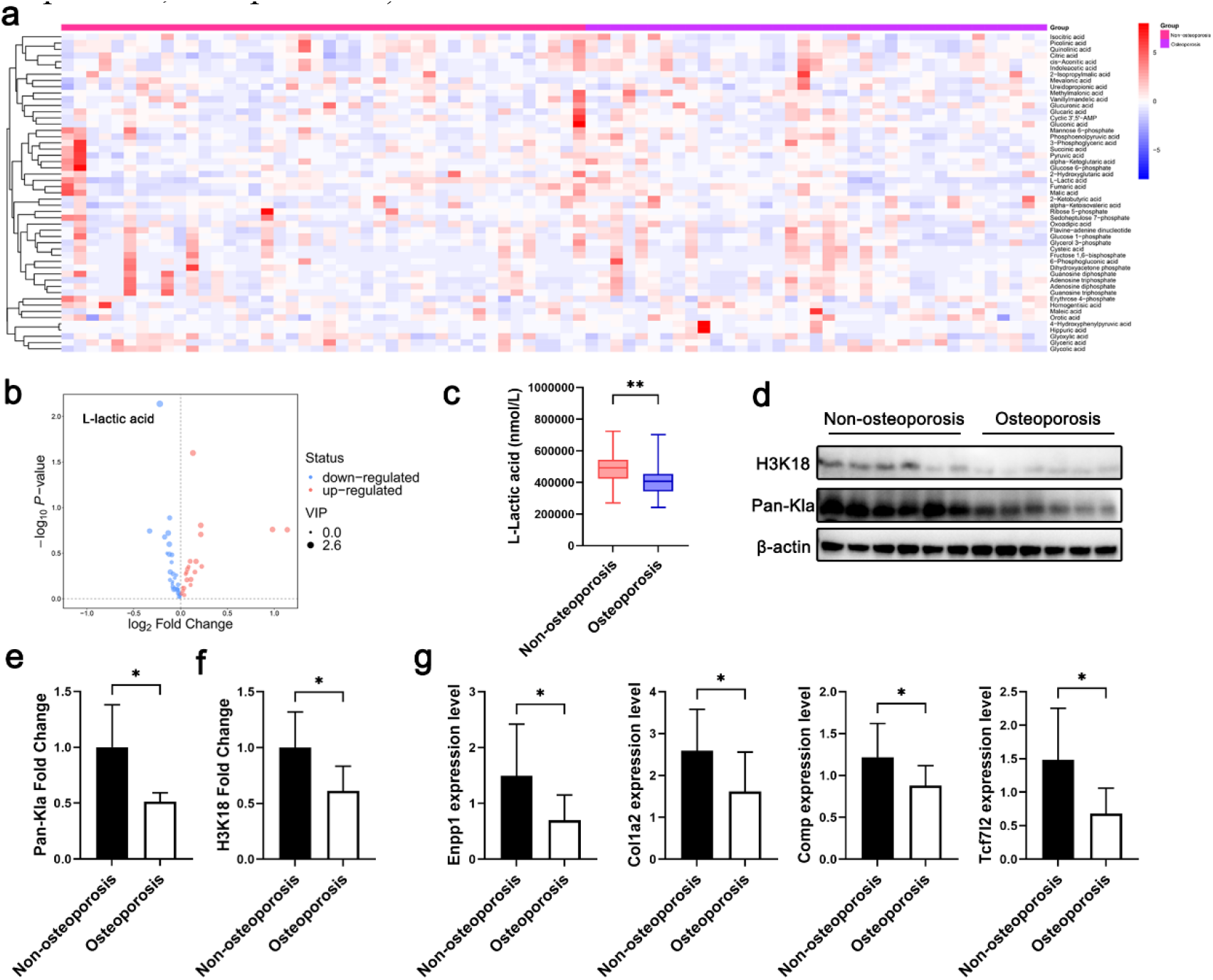
Clinical samples indicate the protective role of lactate in osteoporosis patients. Heatmap of targeted metabolomics of patients’ blood. b) Volcano plot showed the upregulated and downregulated metabolites. c) Lactate was downregulated in osteoporosis group (**p<0.01). d) Histone pan-kla decreased in BMSCs isolated from osteoporosis patients’ bone marrow. e) Quantitative analysis of pan-kla in BMSCs isolated from osteoporosis patients’ bone marrow (*p<0.05). f) Histone H3K18la lactylation decreased in BMSCs isolated from osteoporosis patients’ bone marrow (*p<0.05). g) Quantitative analysis of histone H3K18la lactylation in BMSCs isolated from osteoporosis patients’ bone marrow (*p<0.05). h) qPCR analysis showed decreased levels of COL1A2, COMP, ENPP1, and TCF7L2 in BMSCs isolated from patients’ bone marrow.

## 3. Discussion

Blood vessels transport oxygen, nutrients, growth factors, and metabolites to maintain vial movement[21]. ECs cover the inner wall of the blood vessels, which have recently been found to control osteogenesis[6a]. However, the roles of EC metabolism in osteogenesis and osteoporosis are unclear. The present study found that disruption of ECs glycolysis by specific deletion of PKM2 in ECs suppress osteogenesis and worsen osteoporosis.

Interestingly, the mediator was lactate, a by-product of glycolysis. EC-derived lactate increased BMSCs H3K18la lactylation and upregulated the expression of its target osteogenic genes. Exercise has long been reported to increase osteogenesis and attenuate osteoporosis. In this study, we found that an increased levels of lactate contributed to the therapeutic function of exercise. Importantly, clinical data suggested that high levels of lactate were correlated with high BMD. Decreased H3K18la lactylation and target osteogenic genes were observed in BMSCs isolated from osteoporosis patients. A schematic summarized the mechanism by which ECs glycolysis controls osteogenesis and osteoporosis (**Figure 9**).

**Figure 9.**
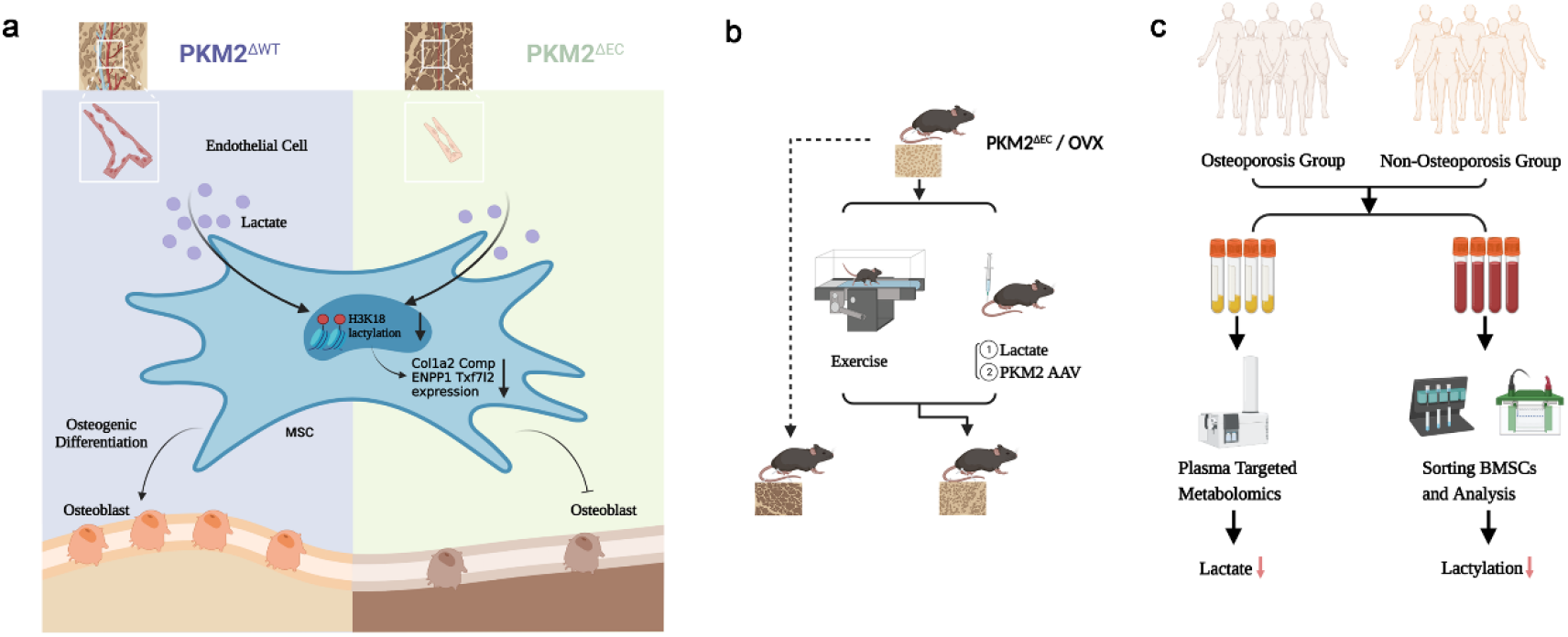
Schematic illustration showing the important role of ECs-derived lactate in osteogenesis and osteoporosis via histone lactylation. a) ECs-derived lactate triggers mesenchymal stem cell histone lactylation to control osteoblast differentiation. b) Exercise, addition of lactate and PKM2 adenovirus improve BMD in OVX mice. c) Lactate levels and BMSCs histone lactylation decreased in osteoporosis group.

Bone angiogenesis plays an important role in bone metabolism, bone remodeling, and bone repair. Vascular endothelial growth factor (VEGF) increases bone blood vessel invasion and regulates growth plate morphogenesis[6b]. Intramembranous ossification requires VEGF for the coupon of angiogenesis and osteogenesis[22]. EC-derived Noggin contributes to bone formation and exogenous Noggin improves the organization of bone vasculature[23]. These studies highlight the importance of angiocrine factors in osteogenesis. The present study found that the density of bone blood vessel decreased in OVX-induced osteoporotic mouse model. ECs are highly glycolytic and generate most of their energy through conversion of glucose to lactate. We found that PKM2, the glycolytic rate-limiting enzyme, is highly expressed in bone blood vessels, and its expression decreased in the OVX model. Using the genetic mice model for PKM2-specific deletion in ECs, we showed that PKM2 controls bone blood vessel formation, which affects osteogenesis. Recent studies reveal that endothelial notch signaling and zinc-finger transcription factor ZEB1 control bone blood vessel and by extension, osteogenesis, in mice[23-24]. However, the mediator of crosstalk between ECs and osteogenic cell lines is not defined. Both Notch signaling and ZEB1 regulate glycolysis, indicating that metabolites may play a role in osteogenesis[25]. These results suggest that the glycolytic metabolite may mediate the crosstalk between ECs and osteogenic cells.

Lactate, previously regarded as a waste product of glycolysis, has received extensive attention as a regulator of multiple molecular processes[26]. Lactate can function as a signaling molecule to affect behavior, shape gut microbiome to alter thermogenesis, and promote PD-1 expression in regulatory T cells to blunt tumor treatment among other function[27]. EC-derived lactate controls adult hippocampal neurogenesis, muscle regeneration from ischemia, and pathological retinopathy[10, 28]. We found that EC-specific deletion of PKM2 resulted in decreased levels of lactate in the blood of mice. Lactate levels were also observed to decrease in the blood of OVX mice. Previous studies indicate that elevated lactate levels may protect against low BMD[29]. This is consistent with our data which showed that osteoporosis patients had relatively low lactate levels. These data highlight the role of lactate in regulation of bone mineral density. The *in vitro* data showed that lactate increased the differentiation of BMSCs toward osteoblast, further revealing the osteogenic function of lactate. Exercise delays the occurrence of osteoporosis; however, the underlying mechanism is unknown[20, 30]. This study found that exercise could increase blood lactate levels and improve osteoporosis in OVX model, thus reinforcing the role of lactate in osteoporosis.

Histone lactylation has been identified as an epigenic modification type for regulating gene transcription[13a]. Recent studies have revealed that histone lactylation occurs widely in human and mouse cells, and controls microglial function, cardiac repair, tumor progression, and inflammation[13b, 16, 31]. During osteoblast differentiation, histone lactylation of the mouse osteoblast precursor cell line increased, suggesting that histone lactylation may participate in osteogenesis[14]. Our data showed that decreased pan-Kla and H3K18la lactylation were observed in BMSCs isolated from PSkm2^ΔEC^ and OVX mice. BMSCs co-cultured with BMECs^Δpkm2^ or treated with conditional cell culture medium from BMECs^Δpkm2^ presented reduced H3K18la lactylation. The joint analysis of CUT & Tag and RNA-seq revealed COL1A2, COMP, ENPP1, and TCF7L2 as the target genes of H3K18la lactylation, which belong to the category of bone mineralization. The expressions of COL1A2, COMP, ENPP1, and TCF7L2 were confirmed using Chip-qPCR. COL1A2 is one of the osteogenic markers related to osteogenesis[32]. COMP mediates bone growth through interaction with extracellular matrix protein 1[33]. ENPP1 is a membrane-bound glycoprotein that controls bone mineralization[34]. TCF7L2 is a key effector of canonical Wnt signaling, which is highly expressed in bones and regulates osteoblast function[35]. These four genes contribute to osteogenesis and bone growth. Importantly, the decreased histone H3K18la lactylation and target genes of COL1A2, COMP, ENPP1, and TCF7L2 were observed in BMSCs isolated from osteoporosis patients. Increasing exogenous lactate levels rescued the inhibited expression of COL1A2, COMP, ENPP1, and TCF7L2 and suppressed H3K18la lactylation in BMSCs isolated from Pkm2^ΔEC^ mice. These results demonstrated that lactate-derived lactylation of H3K18la in BMSCs regulated COL1A2, COMP, ENPP1, and TCF7L2 to control osteoporosis.

## 4. Conclusion

Summarily, we prove that EC glycolysis controls bone blood vessels formation and attenuate osteoporosis through BMSCs H3K18la lactylation. ECs-derived lactate function as angiocrine molecule to mediate the crosstalk between ECs and BMSCs. Exercise help to protect against osteoporosis partially by increasing blood lactate levels. The present study highlights the importance of ECs-derived lactate in osteogenesis and osteoporosis, while also lay the foundation for treatment of osteoporosis by targeting vascular metabolism.

## 5. Experimental section

### Mice

The Pkm2 flox/flox mice were purchased from Cyagen Biosciences. Tg(Cdh5-CreERT2) mice(Strain NO.T014691)were purchased from GemPharmatech (Nanjing, China). Both Pkm2 flox/flox and Cdh5-CreERT2 mice were on a C57BL/6 genetic background. To avoid recombination in the female germline, only Cdh5-cre-positive male mice were used for intercrosses. For constitutive Cre-mediated recombination in ECs, 100mg/kg tamoxifen (Sigma, T5648) in corn oil (Sigma) was intraperitoneally injected to pups every day from P10 to P14 for five consecutive days. Mice were genotyped by PCR performed on genomic DNA. Protocols and primer sequences are accessible upon request. An ovariectomized mouse model was developed as described previously. Summarily, all 8-week old mice were randomly assigned to the OVX and Sham groups. The mice in the OVX group were subjected to bilateral OVX, while those in the Sham group had similar sizes of adipose tissue near the ovaries resected. The ovariectomy operation was performed according to a previously published procedure. Summarily, surgical staff anesthetized mice using isoflurane, removed hair over their dorsum and wiped the skin with povidone-iodine solution. Then the mice were placed in sternal recumbency and an incision was made in the mid-dorsum. Later, operators located each ovary, gently removed the ovary using sterilized fine tweezers, checked for bleeding, and sutured the mice wounds. Finally, the mice were placed in cleaned edges with heating pads until the mice recovered fully. For constitutive Cre-mediated recombination in ECs in OVX or Sham mice, 100 mg/kg tamoxifen in corn oil was administered intraperitoneally to 9-week old mice for 5 consecutive days. All mice were then euthanized 11 weeks later.

For the mice treated with lactate, the mice were subcutaneously injected with lactate every two days after ovariectomy. Moreover, 1.5 g/kg lactate (Sigma) in 0.9% saline whose pH was adjusted to 7.4 was used. The control mice received equal volumes of 0.9 % saline. The treatment of lactate lasted for 8 weeks after ovariectomy.

The mice were subjected to exercise at intervals as previously described[36]. Summarily, each running session was 8 min long at 5 m/min on a rodent treadmill, followed by 10 sessions of high-intensity running (12-16 m/min, 5 min) with 2 min of rest. The high-intensity interval exercise was performed for 5 consecutive days each week and lasted for 8 weeks. The rodent treadmill incline was set at 10°. During the exercise period, electrical stimuli was applied to encourage the mice to run. All mice were kept in alternate dark-light cycles of 12 h at room temperature. All surgeries were performed under sodium pentobarbital anesthesia, and efforts were made to minimize suffering. All animal experiments were conducted in agreement with the NIH Guide for the Care and Use of Laboratory Animals, and with the approval of the Institutional Animal Care and Use Committee at Shanghai Changzheng Hospital.

### Isolation and culture of BMECs

The isolation of BMECs was performed as previously described with minor revision. Summarily, the bone marrow was collected from the tibiae and femurs of mice using sterile liver digestion medium (Gibco) supplemented with 0.1 % DNaseⅠ (Invitrogen). To obtain single BM cell suspension, BM was digested for 30 min at 37°C under shaking condition. BMECs were then sorted using CD31 MicroBeads, mouse (Miltenyi Biotec). Sorted BMECs were seeded on fibronectin (sigma-Aldrich) coated dishes and cultured in endothelial cell medium (ECM, Sciencell) supplemented with endoehlial cell growth supplement, fetal bovine serum, and antibiotic solution (Sciencell) at 37°C 5% CO2.

At first passage, BMECs were sorted using CD31 MicroBeads and plated for culture. The culture medium of BMECs was changed every other day and passaged upon confluency. Passage 2 to 4 of BMECs were used for subsequent experiments.

### CUT & Tag analysis

Hyperactive Universal CUT&Tag Assay Kit for Illumina (Vazyme, TD903) and TruePrep Index Kit V2 for Illumina (Vazyme, TD202) were used for CUT&Tag assay. Summarily, 105 cells were collected and washed once with 500 μl wash buffer, then they were bound to ConA beads for 10 min at room temperature. After that, cells were incubated with first antibody (anti-H3K18La) at 4℃ overnight. The secondary antibodies were added and incubated for 60 min at room temperature. Then cells were washed three times with DIG wash buffer and incubated with 0.04 μM pA/G–Tnp for 60 min at room temperature. Similarly, cells were washed three times with DIG 300 buffer, resuspended in fragmentation buffer and incubated at 37℃ for 1 h. After fragmentation, proteinase K, buffer LB, and DNA extract beads were added to each sample. After incubation at 55℃ for 10 min, beads were rinsed with Buffer WA and Buffer WB. Then DNA were eluted at room temperature for 5 min with 22 μl H2O. CUT&Tag libraries were built using TruePrep Index Kit V2 for Illumina (Vazyme, TD202) following the manufacturer’s protocol.

The libraries were sequenced with an Illumina platform by Shanghai Biotechnology Corporation. The raw reads were preprocessed by filtering out adapters, short-fragment reads, and other low-quality reads. The reads were mapped using Bowtie (version 0.12.8) and called by MACS2 (version 2.2.7.1). The motifs were identified using HOMER software. The data were visualized using Integrative Genomics Viewer. Peaks with fold change ≥2 OR ≤1/2 and p-value<0.05 would be analyzed by R package DiffBind.

### Chip-qPCR

ChIP was performed according to the manufacturer’s instructions (The Agarose ChIP Kit, 26156, ThermoFisher). Summarily, BMSCs were fixed with 1% formaldehyde for 10 min at room temperature and quenched with Glycine Solution (1X) for 5 min. Then, fixed cells were lysed with Lysis Buffer 1 containing protease inhibitors and digested with Micrococcal Nuclease (ChIP Grade) (10U/Μl). The digested chromatin was incubated Dilution Buffer containing primary antibodies (Table *) at 4 ℃ overnight on a vertical roller. Then, adding ChIP Grade Protein A/G Plus Agarose to each IP, centrifuging the column and washing the column with Wash Buffer 1/2/3. Later, eluting IP with 150 μL IP Elution Buffer, 6 μL of 5 M NaCl and 2 μL of 20 mg/mL Proteinase K. Finally, DNA was obtained by DNA Binding Buffer, DNA Clean-Up Column, and DNA Column Wash Buffer following the protocols. Purified DNA levels were quantitatively measured by real-time PCR. The primers were listed in Table S2.

### Immunofluorescence

Femurs was collected from mice and immediately fixed in 4% PFA for 24 h. Decalcification was carried out with 0.5 M EDTA at 4 ℃ with constant slow shaking for 24 h. Then the decalcified femurs were immersed in 20% sucrose and 2% PVP for 24 h. Finally, the bones were embedded and frozen in 8% gelatin using low-profile blades. For immunostaining, bone sections were air-dried, permeabilized for 10 min in 0.2% Triton X-100, and blocked in 5% donkey serum at room temperature for 30 min. Blocked sections were probed with the primary antibodies diluted in 5% donkey serum in PBS for 2 h at room temperature or overnight at 4 ℃. After primary antibody incubation, sections were washed with PBS three times and incubated with appropriate Alexa Fluor-coupled secondary antibodies (1:400) for 1 h at room temperature. Nuclei were counterstained with DAPI. Sections were thoroughly washed with PBS, air-dried, and sealed with nail polish.

### Study population

From February 2021 to November 2022, 182 female patients aged 40 to 55 years who visit the outpatient of department of orthopedics, Changzheng Hospital, were invited to participate in the study. The exclusion criteria included: (1) those who received therapy or medications that may affect BMD, such as hormone replacement therapy or steroid agents; and (2) those who had diseases that may affect bone metabolism, such as fractures or thyroid disorders. In total, 86 participants were included, and all the subjects provided written informed consent. The study protocol was approved by the Committees for Ethical Review of Research involving Human Subjects of Shanghai Changzheng Hospital.

### Statistical Analysis

All data were presented as the means ± standard deviations (SD) from at least three repeated experiments. Statistical analysis was performed using GraphPad Prism (GraphPad software 9.0, GraphPad, Bethesda, MD, USA). Unpaired Student’s t-test was used for the data of two-group analysis, while comparisons between multiple groups were performed by one-way analysis of variance (ANOVA). P values < 0.05 were considered as statistically significant (*P < 0.05; **P < 0.01; ***P < 0.001; ****P < 0.0001).

Additional detailed materials and methods can be found in the Supporting Information.

## Supporting Information

Supporting Information is available from the Wiley Online Library or from the author.

## Acknowledgements

This work was supported by grants from National Natural Science Foundation of China (82172516), Chinese Postdoctoral Science Fund (2020M673673, 2020T130775) and Naval Military Medical University Fund (2021MS10).

## Author Contributions

J. Wu, M. Hu, H. Jiang and J. Ma contributed equally to this work. Funding and study supervision: X. Zhou, C. Wang, T. Lin; Study design: X. Zhou, C. Wang, Y. Gao, J, Wu, M. Hu; Animal studies design: J. Wu, M. Hu, H. Jiang, J. Ma; Data acquisition: J. Wu, M. Hu, H. Jiang, J. Ma, C. Xie, Z. Zhang, X. Zhou, J. Zhao, Z. Tao, Y. Meng, Z. Cai, T. Song, C. Zhang; Data analysis: J. Wu, M. Hu, C. Xie, Y. Gao; Technical support: H. Song, C. Xie; Writing— original draft preparation: J. Wu, M. Hu; writing—review and editing: X. Zhou, C. Wang. All authors have read and agreed to the published version of the manuscript.

## Data Availability Statement

The data that support the findings of this study are available from corresponding author upon reasonable request.

