## Supplemental files for "Endothelial cell-derived lactate triggers mesenchymal stem cell histone lactylation to attenuate osteoporosis"

J. Ma and X. Zhou
Department of Orthopedics, Shanghai General Hospital, Shanghai Jiao Tong University School of Medicine, Shanghai, P.R.China

C. Xie
Department of Neurology, Renji Hospital, Shanghai Jiaotong University School of Medicine

H. Song
Department of Ophthalmology, Changhai Hospital, Naval Medical University, Shanghai, P.R.China

Y. Gao
Senior Department of Orthopedics, The Fourth Medical Center of PLA General Hospital, Beijing, P.R. China

**Keywords:** (vascular metabolism, histone lactylation, lactate, osteoporosis, metabolic reprogramming)

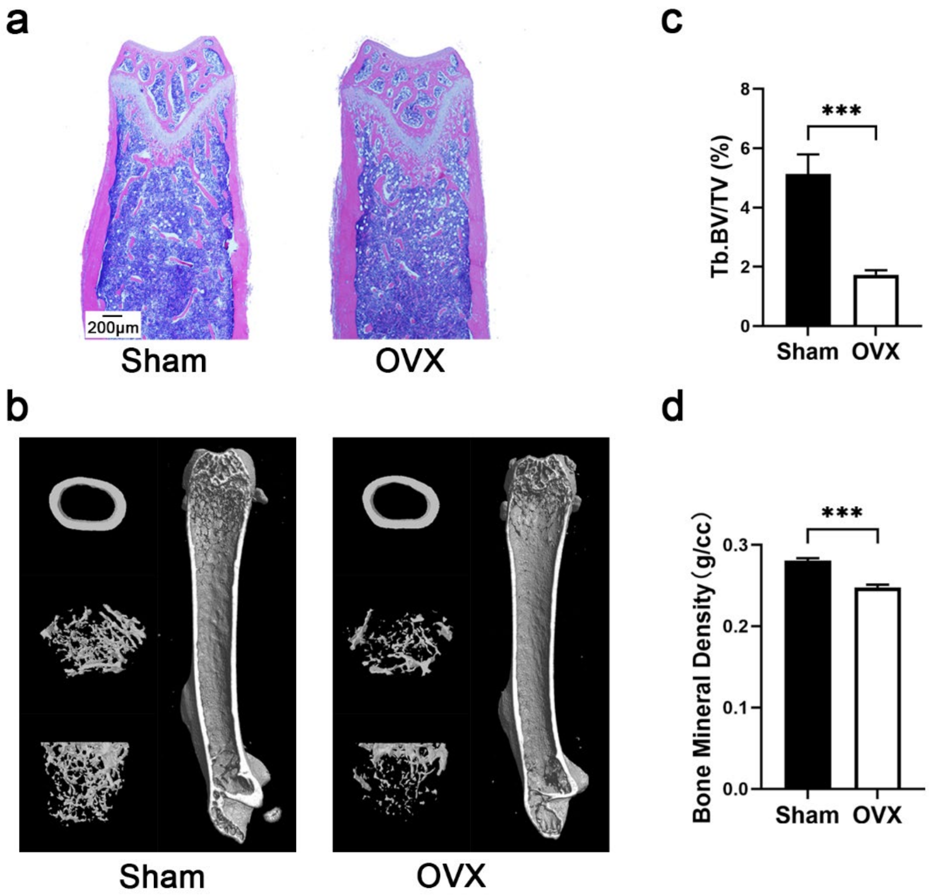


Figure S1. The evaluation of constructed OVX-induced osteoporosis mice. a) Representative images of hematoxylin and eosin staining of distal femur sections. Scale bars 200μ m. b) Representative micro-CT images of the distal femur in OVX mice. c-d) Quantitative analysis of trabecular bone volume per tissue volume (BV/TV) and bone mineral density (BMD) in OVX mice. n=3, (***$p < 0.001$).

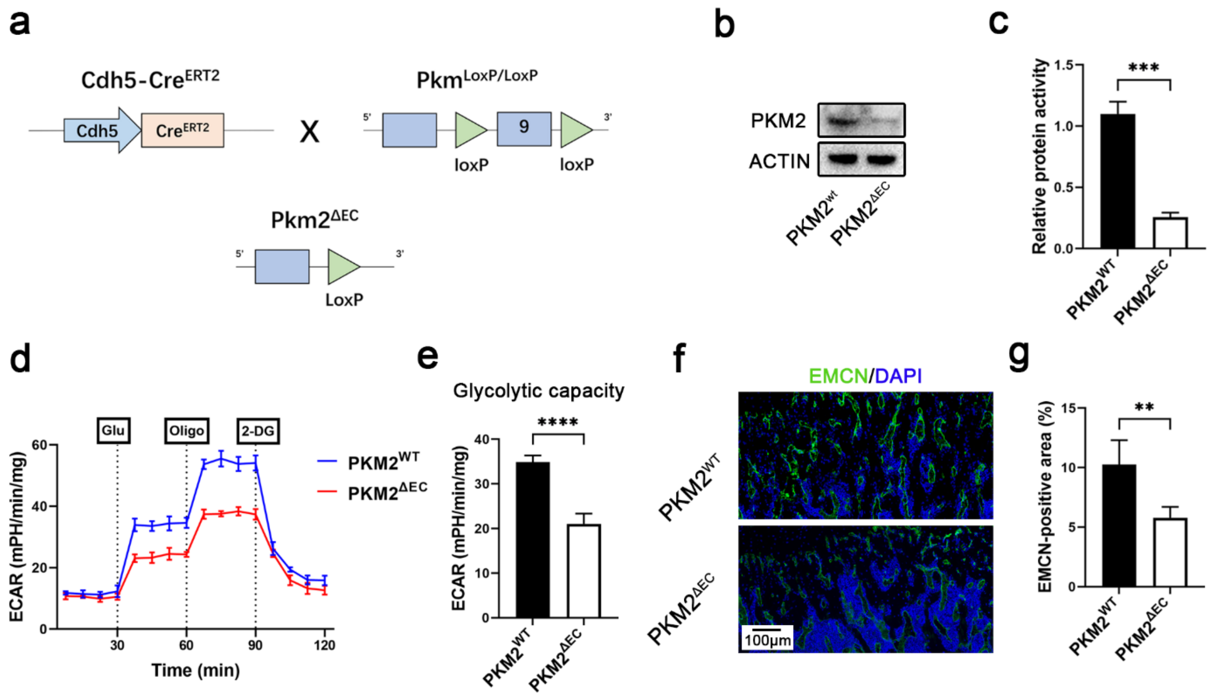


Figure S2. Endothelial PKM2 knockout reduces type H vessels and ECs glycolysis. a) Schematic illustration of breeding for PKM2$^{\Delta EC}$ mice. b) The expression of PKM2 decreased in ECs isolated from Pkm2$^{\Delta EC}$ mice. c) Quantitative analysis of the expression of PKM2. N=3, (***p < 0.001). d) ECAR profile showing glycolytic function of ECs isolated form Pkm2$^{\Delta EC}$ mice. Vertical lines indicate the time of addition of glucose, oligomycin, and 2-DG. e) Quantification of glycolytic function parameters. n=3, (****p < 0. 0001). f) Representative confocal images of femurs stained with EMCN (green) and DAPI (blue). Scale bars, 100 μ m. g) Quantification of CD31 positive vessel area in the BM cavity of the femur sections. n=5, (**p<0.01).

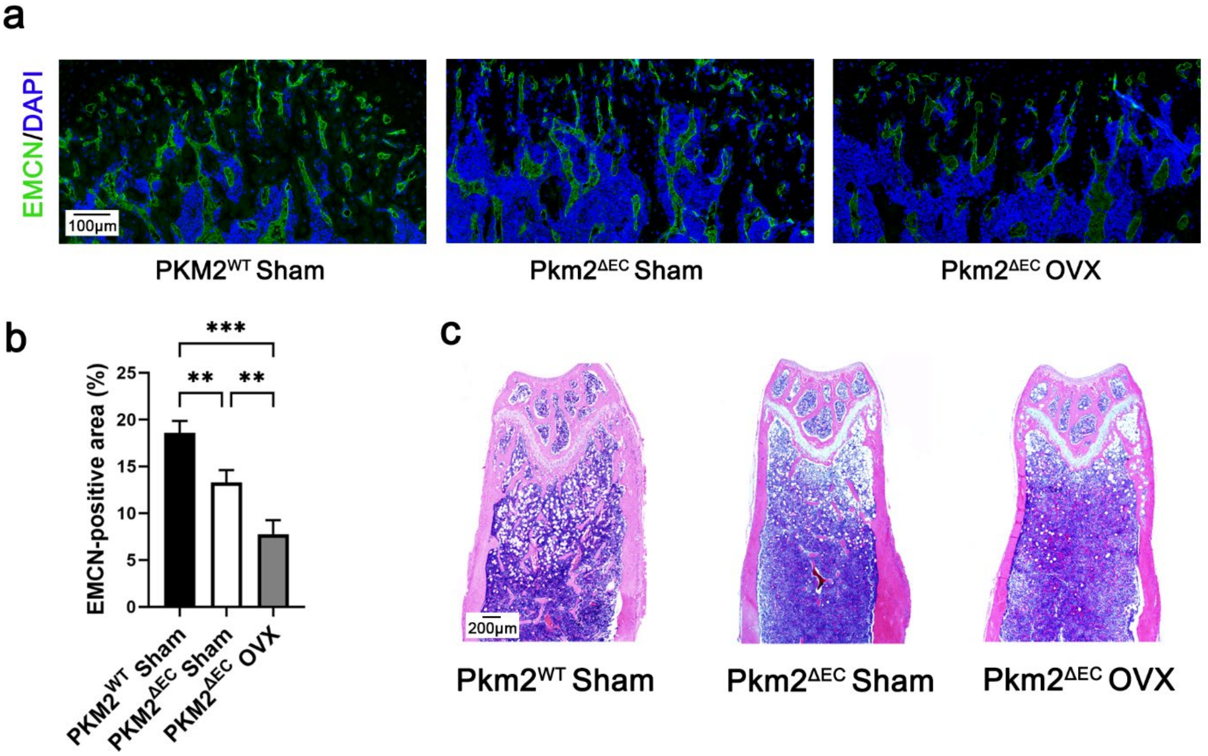


Figure S3. Deletion of endothelial PKM2 worsens osteoporosis. a) Representative confocal images of femurs stained with EMCN (green) and DAPI (blue). Scale bars, 100 μ m. b) Quantification of CD31 positive vessel area in the BM cavity of the femur sections. n=5, (**p<0.01, ***p<0.001). c) Representative images of hematoxylin and eosin staining of distal femur sections. Scale bars 200μ m.

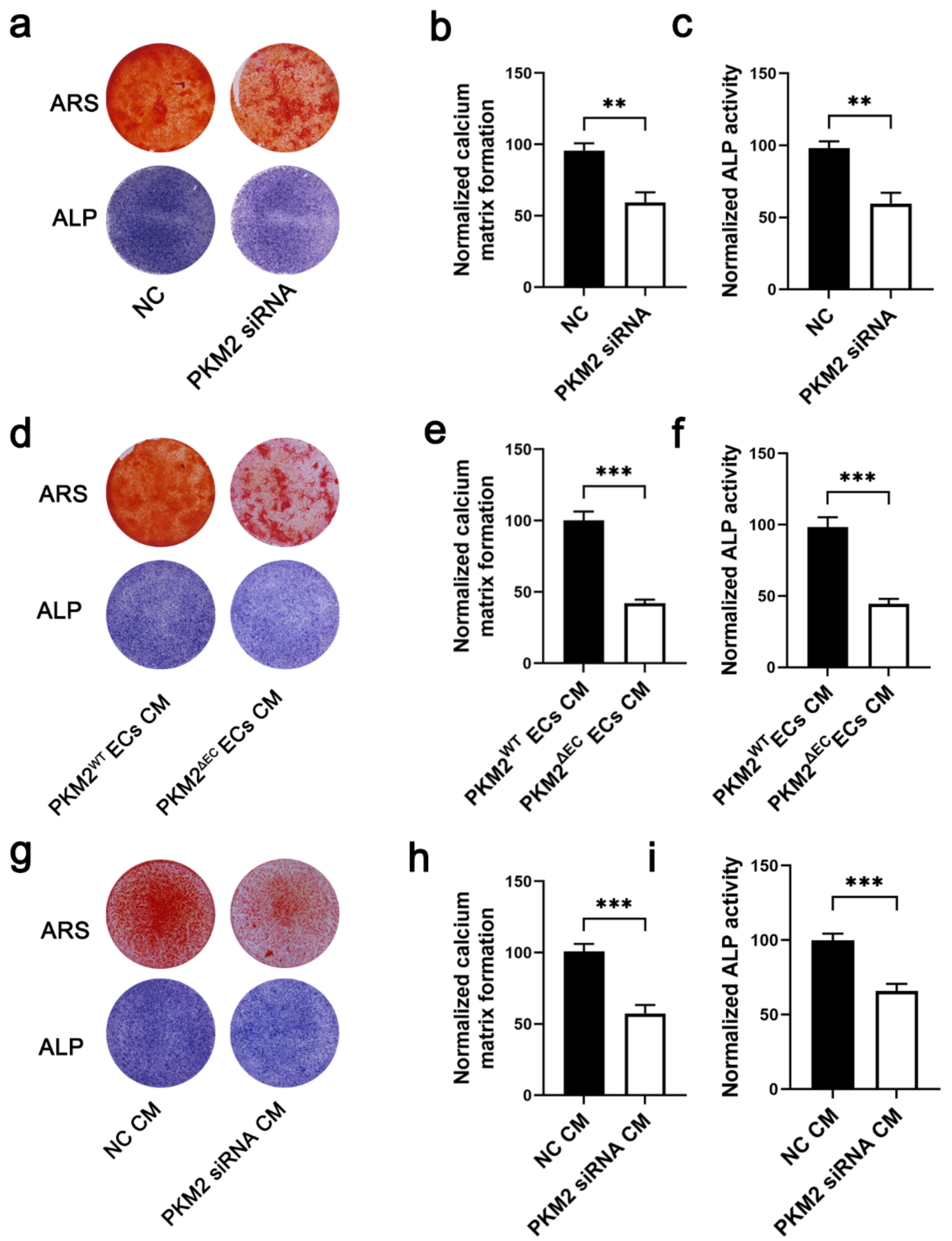


Figure S4. Knock down the expression of endothelial PKM2 suppresses the differentiation of BMSC to osteoblasts. a) Representative images of ARS and ALP staining of BMSCs co-cultured with ECs transfected with PKM2 siRNAs. b, c) Quantitative analysis of ARS and ALP staining of BMSCs co-cultured with ECs transfected with PKM2 siRNAs. n=4, (**p<0.01). d) Representative images of ARS and ALP staining of BMSCs cultured with conditional medium from BMECs$^{\Delta pkm2}$ and BMECs$^{wt}$. e, f) Quantitative analysis of ARS and ALP staining of BMSCs cultured with conditional medium from BMECs$^{\Delta pkm2}$ and BMECs$^{wt}$. n=4, (***p<0.001). g) Representative images of ARS and ALP staining of BMSCs treated with conditional cell culture medium from ECs transfected with PKM2 siRNAs. h, i)

Quantitative analysis of ARS and ALP staining of BMSCs treated with conditional cell culture medium from ECs transfected with PKM2 siRNAs. n=4, (***p<0.001).

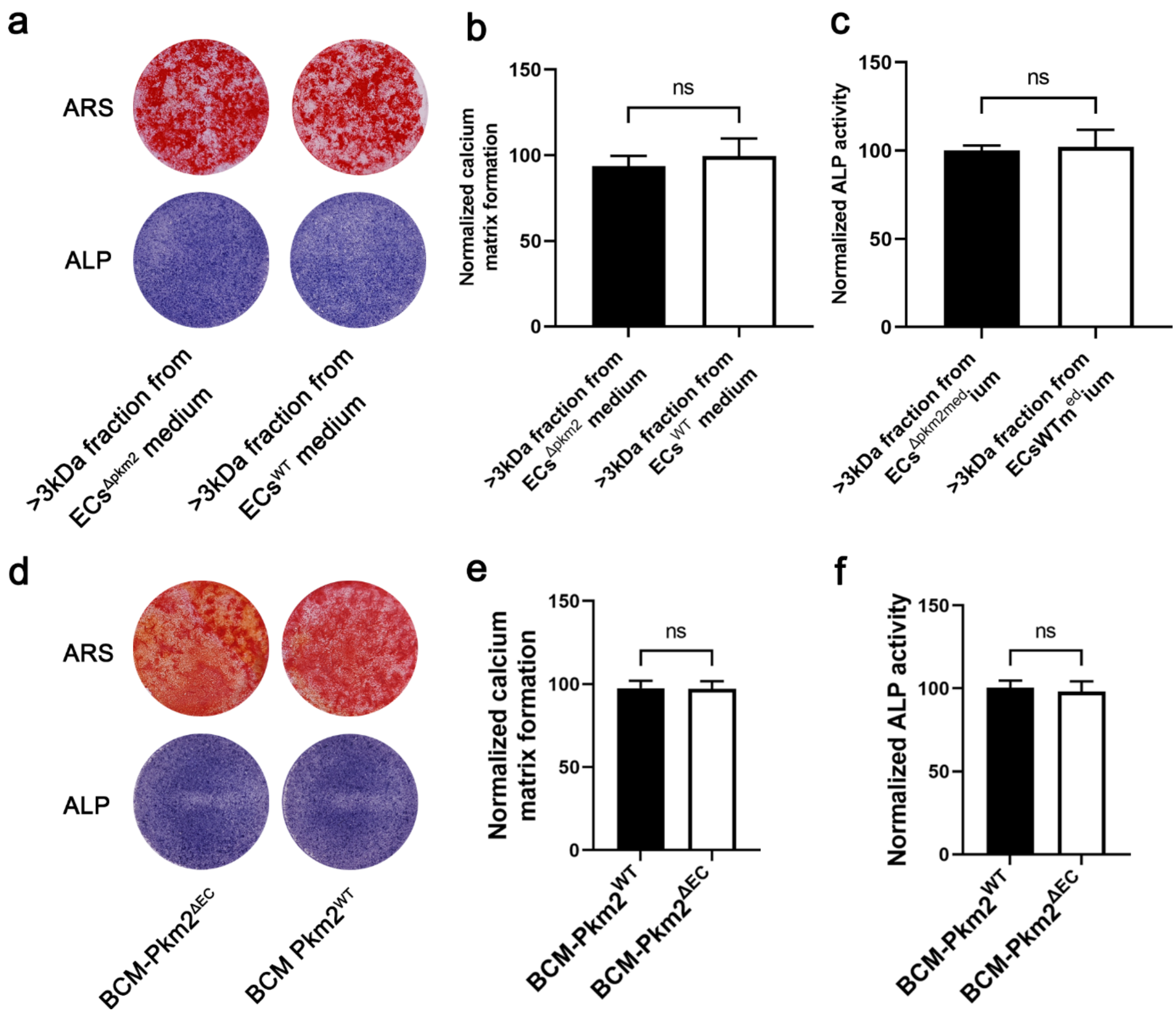


Figure S5. Large fractions (>3kDa) of conditional medium does not affect the differentiation of BMSC. a) Representative images of ARS and ALP staining of BMSCs cultured with conditional medium fractions that >3kDa. b, c) Quantitative analysis of ARS and ALP staining of BMSCs cultured with conditional medium fractions that >3kDa. n=4. d) Representative images of ARS and ALP staining of BMSCs cultured with boiled conditional medium from BMECs$^{\Delta pkm2}$ and BMECs$^{wt}$. e, f) Quantitative analysis of ARS and ALP staining of BMSCs cultured with boiled conditional medium from BMECs$^{\Delta pkm2}$ and BMECs$^{wt}$. n=4.

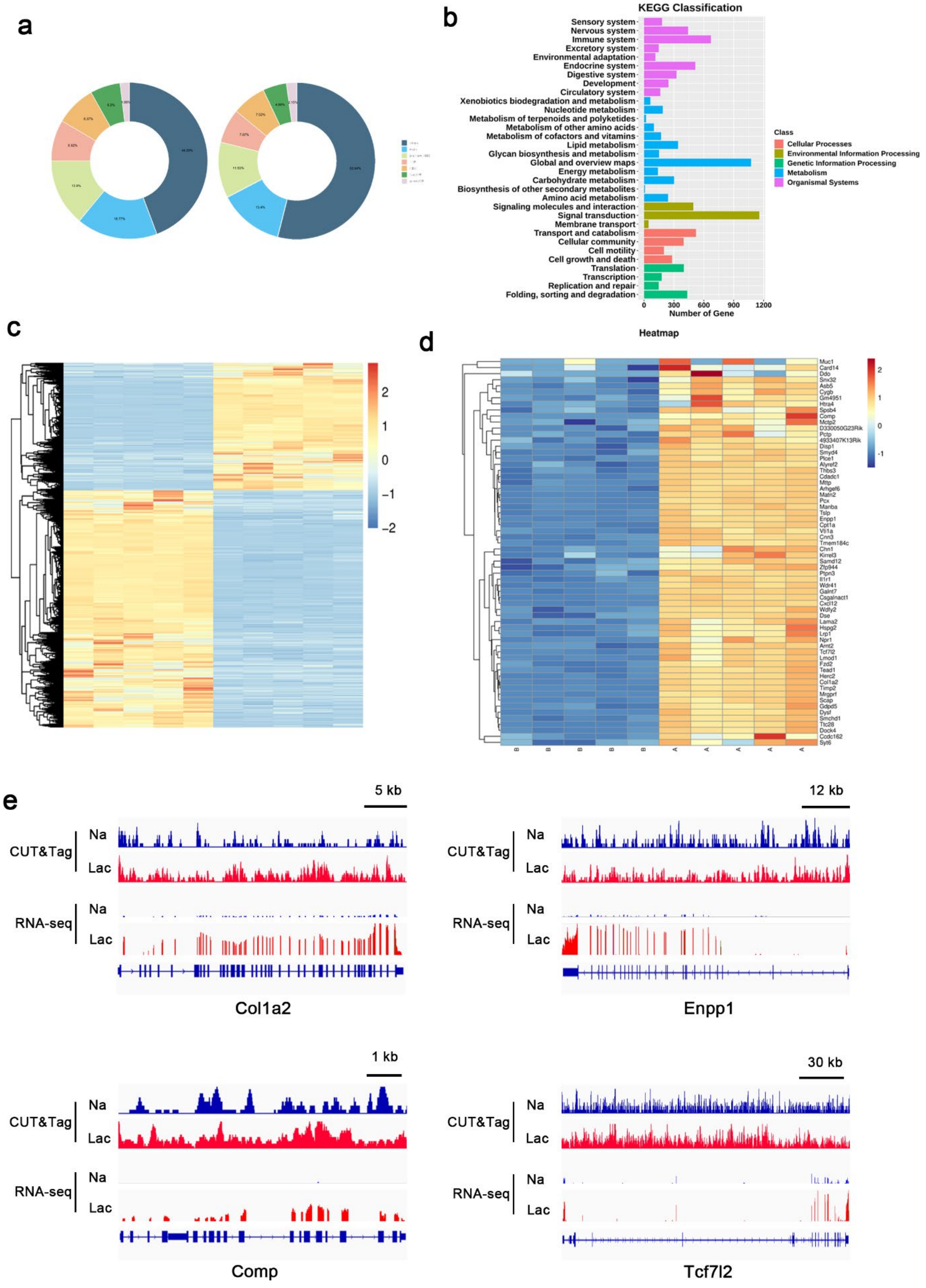


Figure S6. Bioinformatics analysis of CUT&Tag and RNA-seq data. a) b) c) Heatmap of differential expressed genes of RNA-seq. d) Heatmap of

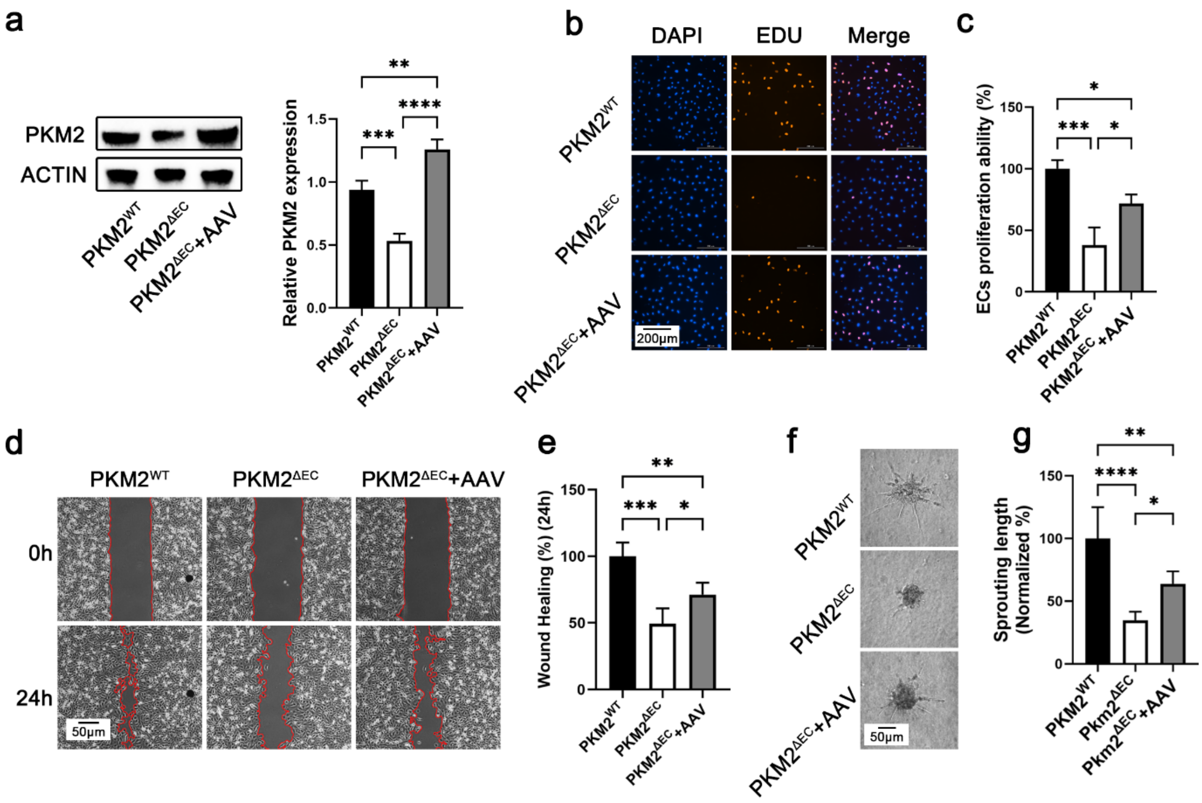


Figure S7. Overexpression of PKM2 in ECs improve the angiogenic function. a) The expression of PKM2 was rescued by AAV in ECs isolated from Pkm2$^{\Delta EC}$ mice. Quantitative analysis of the expression of PKM2. n=3, (**p<0.01, ***p<0.001, ****p<0.0001). b) Cell proliferation determined by EdU staining. EdU (Red), DAPI (Blue), (scale bar 200 μm). c) Statistical result of cell proliferation. n=4, (**p<0.01). d) Representative images of migrated cells (scale bar 50 μm). e) Statistical result of cell migration. n=4, (*p<0.05, **p<0.01, ***p<0.001). f) Representative images of cell sprouting (scale bar 50 μm). h) Statistical result of cell sprouting. n=4, (*p<0.05, **p<0.01, ****p<0.0001).

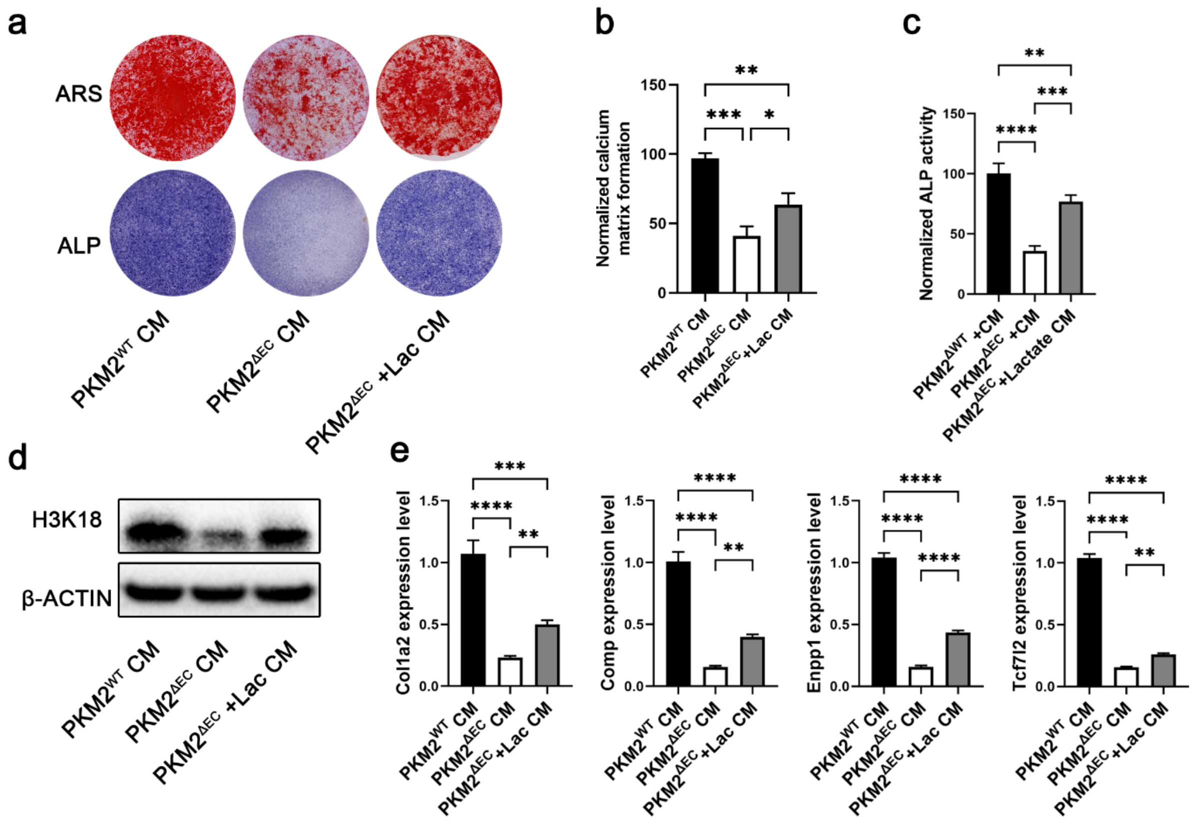


Figure S8. Lactate increases the lactylation and the differentiation of BMSC. a) Representative images of ARS and ALP staining of BMSCs isolated from PKM2$^{\Delta EC}$ mice and treated with lactate. b, c) Quantitative analysis of ARS and ALP staining of BMSCs isolated from PKM2$^{\Delta EC}$ mice and treated with lactate. n=4, (*p<0.05, **p<0.01, ***p<0.001). d) The expression of K3K18la was rescued by exogenous lactate. e) qPCR analysis of COL1A2, COMP, ENPP1, and TCF7L2 in BMSCs cultured with medium from ECs supplemented with exogenous lactate. n=3, (**p<0.01, ***p<0.001, ****p<0.0001).

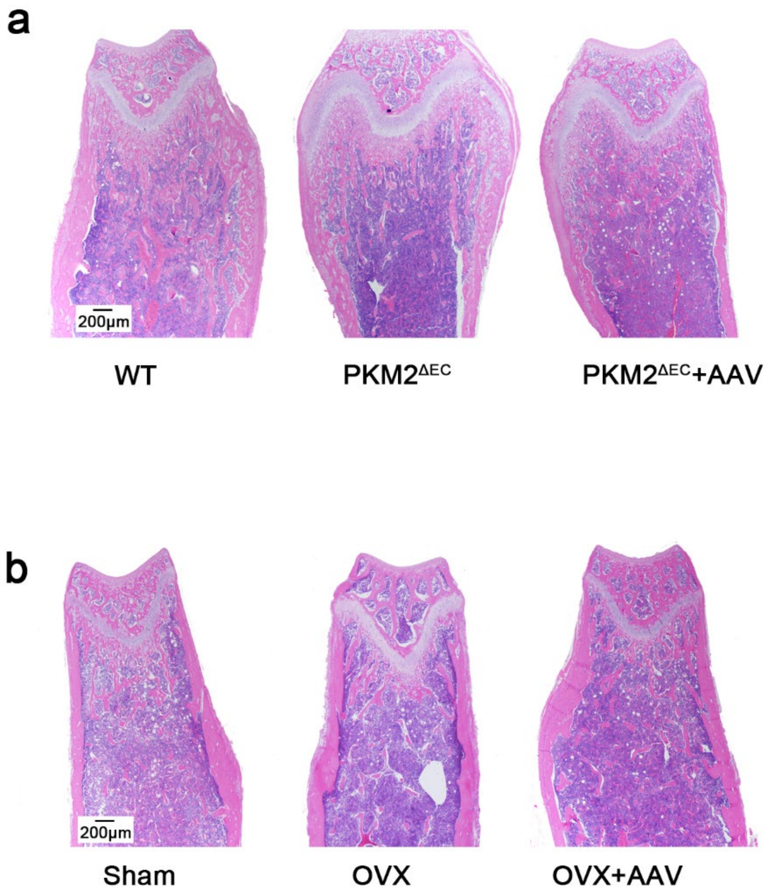


Figure S9. PKM2 adenovirus improve osteogenesis and osteoporosis in PKM2$^{\Delta EC}$ mice. a) Representative images of hematoxylin and eosin staining of distal femur sections in PKM2$^{\Delta EC}$ mice treated with PKM2 adenovirus. Scale bars 200μ m. b) Representative images of hematoxylin and eosin staining of distal femur sections in OVX PKM2$^{\Delta EC}$ mice treated with PKM2 adenovirus. Scale bars 200μ m.

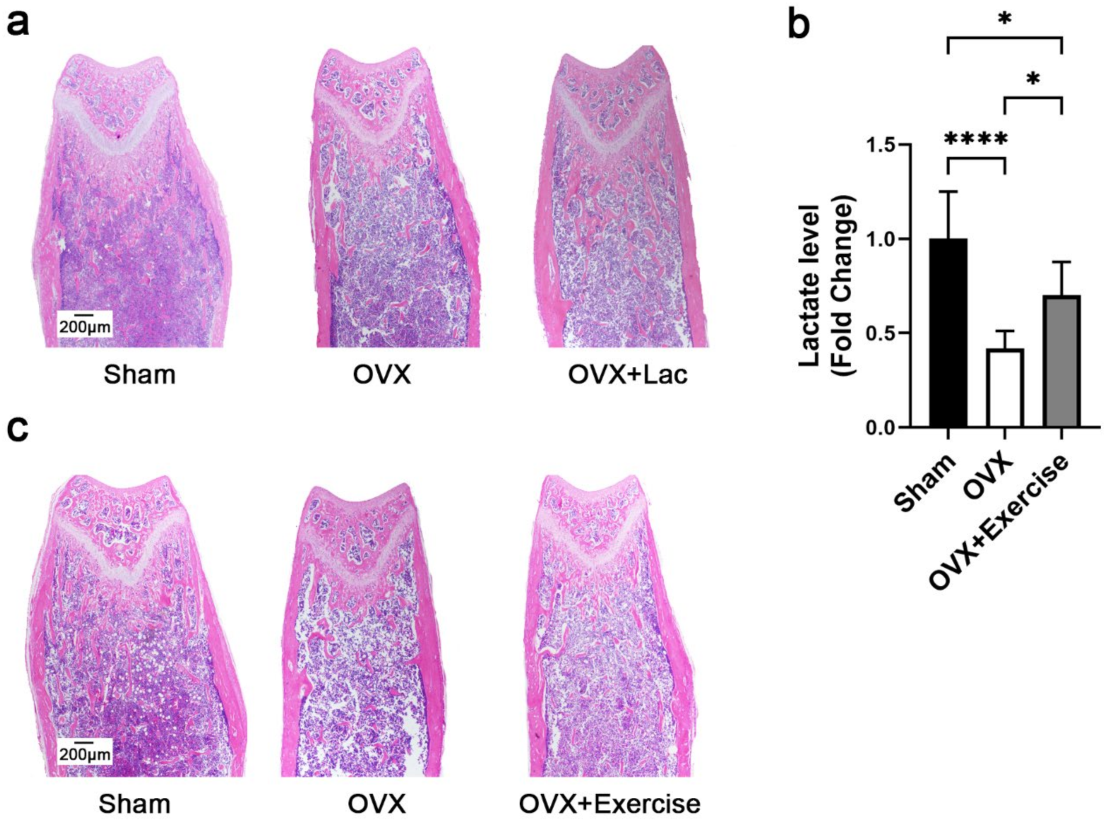


Figure S10. Exogenous lactate and exercise improve osteoporosis. a) Representative images of hematoxylin and eosin staining of distal femur sections in OVX mice treated with lactate. Scale bars 200μ m. b) high-intensity interval exercise increased blood lactate levels in OVX mice. c) Representative images of hematoxylin and eosin staining of distal femur sections in OVX mice with intense exercise. Scale bars 200μ m.

**Supplementary Table 1**

Baseline characteristics of study participants

| | Non-osteoporosis | Osteoporosis | ALL |
|---|---|---|---|
| **Age** | 47.74±4.31 | 47.84±3.64 | 47.78±3.99 |
| **FBG** | 5.70±0.80 | 5.61±0.71 | 5.65±0.65 |
| **BMI, kg/m$^2$** | 23.76±4.14 | 23.44±4.06 | 23.61±4.08 |
| **BMD T (Lumbar spine )** | -0.62±0.23 | -2.75±0.22 | -1.62±1.10 |

FBG: fasting blood glucose

**Supplementary Table 2**

Primers used for RT-qPCR

| Primers name | Sequences (5‘ to 3‘) |
|---|---|
| Enpp1 F | GAGCTCGAGTCACCAGCC |
| Enpp1 R | CACATACTGACAAAACCAGCGA |
| Comp F | AGTTGGCTACATCAGGGTGC |
| Comp R | CCGTGTCCAACACCACATTG |
| Tcf7l2 F | GGGAAACCAACGAACACAGC |
| Tcf7l2 R | GGGAAACCAACGAACACAGC |
| Col1a2 F | TAAAGAAGGCCCTGTGGGTC |
| Col1a2 R | ATCCGATGTTGCCAGCTTCA |
| β-Actin F | CGTTGACATCCGTAAAGACC |
| β-Actin R | AACAGTCCGCCTAGAAGCAC |

**Supplementary Table 3**

Primers used for ChIP-qPCR

| Primers name | Sequences (5‘ to 3‘) |
|---|---|
| Enpp1 F | AGCAAGGAAGGAGCCTCTTG |
| Enpp1 R | AGCCTTGTGATATTTGGCTCAAG |
| Comp F | CAGTCCAGTGAGAAGAGAACCC |
| Comp R | CTTCCCCTTTTGCACTGTTGAG |
| Tcf7l2 F | CTTGTCATCAGGGGACGATCTA |
| Tcf7l2 R | TGTAGACAAACACCTTCAGAC |
| Col1a2 F | AGCTTGGGATGTTAAGGAAGA |
| Col1a2 R | GTTGATGAGGGGAGCAGGCAT |

### Supplementary Experimental Section

*Isolation of primary BMSCs from mice:* The protocol for isolating BMSCs from mics was reported previously. Summarily, the mice were euthanized, then femurs and tibias were cleaned thoroughly to remove muscle and connective tissue. Then small cuts were made at the ends of bones using sterile scissors and bone marrow was blown out using sterile syringes. Later, bone marrow cells were seeded on 10 cm dishes for pair of mice. After culturing for 48 h, the medium was changed with complete medium (α-MEM with 20% FBS and 1% Penicillin/Streptomycin) to remove non-adherent cells and were changed every 3 days. Cells at passages 2–4 were employed for the following in vitro experiments.

*Isolation of primary BMSCs from patients:* From February 2021 to November 2022, twelve female patients aged 40 to 55 years who underwent surgery at the department of orthopedics, Changzheng Hospital, participated in the study. Bone marrow was collected from patients, who provided informed consent, during the surgery. BMSCs were isolated according to a previously published protocol.
Summarily, bone marrow mononuclear cells (BMMNCs) were isolated and purified from the bone marrow samples by density gradient centrifugation methods on Ficoll (P4350, Solarbio). The bone marrow samples were attenuated with sterile PBS (1:2), layered onto equal volume Ficoll, and the mixtures were isolated by centrifugation (440 g, 20 min, 4℃). The BMMNCs layer would be transferred into a new 15 ml centrifuge tube and washed twice with PBS. The red blood cells were lysed using RBC lysis buffer. Later, CD271 magnetic MicroBeads Kits (human) (130-099-023, Miltenyi Biotec) were used to purify the remaining cells of each sample. The samples were re-suspended in 60 μl buffer, 20 μl of FcR Bloking Buffer, and 20 μl of CD271 MicroBeads per $10^7$ cells. After incubation at 4°C for 15 min, the samples were washed using a 2 ml buffer and re-suspended in 500 μl of buffer. Finally, the cells were applied to the LS columns and the labeled $CD271^+$ cells were collected for subsequent experiments.

*Osteogenic differentiation:* Primary BMSCs were seeded into plates in complete medium. After the cells reached 80–90% confluence, the medium was changed to an osteogenic differentiation medium containing α-MEM, 10% FBS, β-glycerophosphate (20 mM), dexamethasone (10 nM), and ascorbic acid (50 mM). The differentiation medium was replaced every 2 days.

*Preparation of BMECs-CM:* BMECs sorting from $Pkm2^{\Delta EC}$ or BMECs treated with PKM2 siRNAs were cultured with fresh ECM when they reached approximately 40% confluence at 37°C, 5% CO2. Supernatants were collected on day 3 and transferred to the 50 mL centrifuge tube. Conditional medium from BMECs was then centrifuged at 2000 g for 10 min to remove cellular debris. To acquire sterile BMECs-CM, conditional medium was filtered using 0.22 μm filter with a 50 ml syringe barrel. Amicon Ultra-15 centrifugal filters were used to separate fractions that were > 3 kDa and < 3 kDa. The BMECs-CM fraction that was larger than 3 kDa remained above the filter and the smaller fraction passed through to the lower chamber. The larger fraction was resuspended in ECM for further usage.

*BMECs and BMSCs co-culture assay:* The co-culture of BMECs and BMSCs was achieved using the transwell (pore size = 0.4 μm) system. Approximately 1×104 BMECs were seeded in the upper chambers of transwell and 2×104 BMSCs were seeded in the well of the 24-well plate. The upper chamber was filled with ECM to culture BMECs and the mesenchymal stem cell osteogenic differentiation medium was used to culture BMSCs. The culture mediums were changed to fresh mediums every 3 days. After co-culturing for 10 days, BMSCs were rinsed with PBS, and processed for subsequent experiments.

*RNA-seq analysis:* Total RNA was isolated from BMSCs using TRIzol regent. The RNA was extracted in 1.5 ml cryopreservation tubes in liquid nitrogen. After RNA extraction, RNA quality and integrity was detected by NanoDrop ND-1000 (NanoDrop, USA) and Bioanalyzer 2100 (Agilent, USA). Subsequently, Dynabeads Oligo (dT) (Thermo Fisher, cat.25-61005, USA) was used to purify RNA. cDNA libraries were generated using NEBNextR Magnesium RNA Fragmentation Module kit (BioLabs, cat.E6150S, USA) and SuperScriptTM II Reverse Transcriptase, (Invitrogen, cat.1896649, USA). The RNA libraries were sequenced on the illumina NovaseqTM 6000 platform by LC Bio Technology CO, Ltd (Hangzhou, China) with the PE150 model. R package DESeq2 was used to conduct differentially expressed genes (DEG) analysis, which was identified with fold change > 2 and $p < 0.05$.

*Quantitative real-time RT-PCR:* Total RNA was extracted and purified with the RNA-Quick Purification Kit (RN001, YiShan Biotech) and reverse transcribed into cDNA using the PrimeScript RT Master Mix (RR036A, TaKaRa) according to the manufacturer's instructions. TB Green Premix Ex Taq II (RR820A, TaKaRa) and primer mixtures were used for real-time RT–PCR. Gene expression levels were normalized relative to the expression of β-actin. The ΔΔCt method was used for the quantification of gene expression. The primers were listed in Table S3.

*Western Blot:* Total proteins were extracted with RIPA with PMSF. The proteins were separated by SDS-PAGE and transferred to PVDF membranes (Millipore). Next the membranes were blocked with 3% BSA and incubated with the primary antibodies at 4℃ overnight. Then the membranes were incubated with secondary antibodies for 60 min at room temperature. Finally, the signals were visualized using a Bio-RAD Image software.

*ALP staining:* BMSCs were incubated in osteogenic differentiation medium for 7 days. The cells were washed twice with PBS and fixed for 30 min in 4% PFA at room temperature. Then the ALP staining was performed with the kit (C3206, Beyotime) at 37℃ for 30 min. ALP activity was quantified using the Alkaline Phosphatase Assay Kit (P0321S, Beyotime)

*Alizarin Red S staining:* BMSCs were plated in 12-well plates cultured in complete medium. The complete medium was changed to osteogenic differentiation medium after the cell confluence reached 90%. Cells were washed in double distilled water twice and fixed in 4% PFA at room temperature. Then the cells were stained with 2% Alizarin Red S solution (G1450, Solarbio), washed twice with double distilled water, and photographed to detect calcification. Finally, 1 ml of 0.1 M NaOH solution was added and semi-quantitative analysis performed by detecting absorbance at 480 nm.

*Micro-CT analysis:* The femurs were fixed in 4% PFA-PBS and analyzed using high-resolution micro-CT () and software. The metaphysis of the tibia was scanned and metaphyseal parameters such as Tb.BV/TV and BMD were measured.

*Plasma lactate measurements:* Blood samples were acquired from the mice saphenous vein puncture as previously described [saphenous vein puncture for blood sampling of the mouse, rat, gerbil, guinea pig, ferret and mink] and mixed with 0.5 M EDTA (1:500). Then, the samples were centrifuged at 3000 rpm for 20 min and supernatant was collected carefully for the subsequent analysis. The concentrations of plasma lactate in mice were measured using the Lactic Acid Test Kit (A019-2-1, Nanjing Jiancheng Bioengineering Institute) according to the manufacturer's instructions. The test tube contained 20 μL sample plasma, 1 ml Enzyme Working Buffer, and 20 μL Chromogenic Solution and the control tube of each sample contained 20 μL standard Lactic Acid Solution, 1 ml Enzyme Working Buffer, and 20 μL Chromogenic Solution. After incubating at 37 °C for 10 min, Stop Solution was added to each tube. Finally, the absorbance value of each sample was measured at 530 nm. The lactic acid concentration was analyzed depending on the formula according to the manufacturer's instructions.

*Scratch-wound assay:* Summarily, 1×105 cells were plated in a 2-well Ibidi chamber plate to form a confluent monolayer. Afterwards, the insert was pulled out and cells were rinsed with PBS three times to remove the suspended cells, and then cultured with the original medium. Images were obtained at 0 h and 16 h using a microscope (IX81, Olympus). All images were processed using ImageJ software.

*Spheroid assay:* Spheroid sprouting assay was performed as previously described. Summarily, BMECs were digested and counted using a hemacytometer and cells were resuspended into medium to acquire a density of 20000 cells/mL. The cells were mixed with 1 ml of methocel stock solution and transferred to a sterile multichannel pipette reservoir. Then 25 μl of the solution was pipetted onto a 10 cm square petri dish. Then the dish was inverted and incubated in cell culture incubator for 24 h. The spheroids were gently washed with 10 ml PBS and transferred into a 15 ml conical tube. Spheroids were centrifuged at 200 g for 5 min and then resuspended in 2 ml of methanol containing 20% FBS. Then 4 ml collagen solution was mixed with 0.5 ml 10× Medium 199, and the pH value was adjusted by adding sterile ice-cold 0.2 N NaOH. Furthermore, 2 ml of the collagen/Medium 199 solution was mixed with methanol solution containing spheroids. Then 1 ml of the spheroid-collagen solution was added per well on a 24-well plate, which was incubated at 37 °C for 30 min to allow the collagen to polymerize. Spheroids were simulated with 200 μL of ECM or ECM containing BREs for 24 h in a humidified incubator (37 °C, 5% CO2). Images were captured using a microscope (IX81, Olympus) and sprouting numbers or vascular length were calculated using ImageJ software.

*EdU proliferation assay:* Cellular proliferation was assessed using 5-ethynyl-2-deoxyuridine (EdU) according to the manufacture's instruction. Cells were co-cultured with EdU working solution (1:1000) for 2 h, followed by fixation with 4% paraformaldehyde for 30 min. Glycine was then added to wash the cells for 5 min, followed by two washes with 0.3% Trion X-100. Subsequently, the cells were incubated with Apollo fluorescent azide for 30 min at room temperature in the dark, followed by three washes with 0.3% Trion X-100. DAPI was incubated for another 30 min, and the cells were washed three times with PBS. The proliferation was calculated as the number of EdU-positive cells/number of DAPI-stained cells. Images were acquired using a fluorescence microscope (IX81, Olympus), and cell counting was performed using the ImageJ software.
*Adeno-associated virus production and administration:* The virus packaging was supported by OBiO Technology Corp, Ltd (Shanghai). The serotype 9 adeno associated virus (AAV9) was used as the vector to carry mice Pkm2 cDNA. AAV9-empty was used as control. The titer used was $2\times10^{11}$ virus genome/mouse. In order to rescue the expression of PKM2 in endothelium, AAV9 was employed to manipulate the expression of PKM2 *in vivo* via tail vein injection. The expression of PKM2 and relevant experiments were performed 6 weeks later after AAV9 injection.

*Targeted metabolomics:* Targeted metabolomics profiling was conducted by Shanghai iProteome Biotechnology Co., Ltd. The plasma samples were collected in Changzheng hospital. In addition, 100 μL of each sample was precisely mixed with 300 μL of methanol (precooled at -40 °C) and vortexed for 30 s. After sonication in an ice-water bath for 15 min and incubation at -40 °C for 60 min, the samples were centrifuged for 15 min (12000 rpm, 4 °C) and 320 μL aliquot of the clear supernatant was collected and dried by spin. Finally, 160 μL of ultrapure water was added to dissolve the dried samples, which were then transferred to inserts in injection vials for HPIC-MS/MS analysis.

The cell samples were collected in a 1.5 ml sterile centrifugal tube and the supernatant was discarded after centrifugation. Then the cell samples were frozen in liquid nitrogen for 15 s. Later, 300 μL of water was added to each sample, and the samples were vortexed for 30 s. The samples were subjected to freeze-thaw three times in liquid nitrogen, vortexed for 30 s, and sonicated for 15 min. Then, 250 μL aliquot of the clear supernatant was mixed with 750 μL aliquot of methanol, containing isotopically-labelled internal standard, precooled at -40 °C. After incubation at -40 °C for 60 min, the samples were centrifuged for 15 min (12000 rpm, 4 °C) and 900 μL clear supernatant was collected and dried by spin. Finally, 180 μL of water was added to the dried residue to dissolve the dried samples, which were then transferred to inserts in injection vials for HPIC-MS/MS analysis.

Multivariate analysis was performed using SIMCA16.0.2 software (Sartorius Stedim Data Analytics AB, Umea, Sweden). Data were scaled and logarithmic transformed to minimize the impact of both noise and high variance of the variables. Principle component analysis (PCA) was performed to visualize the distribution and grouping of the samples. Moreover, 95% confidence interval in the PCA score plot was used as the threshold to identify potential outliers in the dataset.

*Seahorse flux analysis:* To allow comparison between experiments, data are presented as the extracellular acidification rate (ECAR) in mpH/min/μg protein. BMECs were seeded on XF-96 plates with 15000 cells/well in 200 μL of complete medium 48 h before the experiment. Cells were washed and the glucose medium was removed to perform ECAR analysis. Then, the ECAR was measured following the sequential addition of glucose (20 mM), FCCP (2 μM) and antimycin (25 mM). Later, ECAR was automatically calculated using the Seahorse XF-24 analyzer (Seahorse Bioscience, USA).
